# Quantifying the heterogeneity of macromolecular machines by mass photometry

**DOI:** 10.1101/864553

**Authors:** Adar Sonn-Segev, Katarina Belacic, Tatyana Bodrug, Gavin Young, Ryan T. VanderLinden, Brenda A. Schulman, Johannes Schimpf, Thorsten Friedrich, Phat Vinh Dip, Thomas U. Schwartz, Benedikt Bauer, Jan-Michael Peters, Weston B. Struwe, Justin L. P. Benesch, Nicholas G. Brown, David Haselbach, Philipp Kukura

## Abstract

Sample purity is central to *in vitro* studies of protein function and regulation, as well as to the efficiency and success of structural studies requiring crystallization or computational alignment of individual molecules. Here, we show that mass photometry (MP) accurately reports on sample heterogeneity using minimal volumes with molecular resolution within minutes. We benchmark our approach by negative stain electron microscopy (nsEM), including workflows involving chemical crosslinking and multi-step purification of a multi-subunit ubiquitin ligase. When applied to proteasome stability, we detect and quantify assemblies invisible to nsEM. Our results illustrate the unique advantages of MP for rapid sample characterization, prioritization and optimization.

---

Biomolecular structure is frequently affected by non-covalent interactions resulting in mixtures of species even for highly purified samples under physiological conditions^1^. Most proteins perform their function only in specific assemblies ranging from monomeric species for simple binding^2^, to large, heterooligomeric molecular machines^3^. At the same time, one of the major remaining bottlenecks to routine and high-throughput structure determination is sample homogeneity,^4^ which is of immense importance for both cryo-electron microscopy^5,6^ and x-ray crystallography^7^.

Much effort has therefore been aimed at developing and optimizing techniques capable of accurately reporting on sample heterogeneity including SDS-PAGE, size exclusion chromatography (SEC) and dynamic light scattering (DLS)^8,9^. SDS-PAGE reveals sample size, but not stoichiometry or interactions. SEC reports on stokes radii, not the actual molecular weight, and DLS has limited mass accuracy and resolution. For protein complexes, negative staining EM (nsEM) thus remains the standard method for evaluating sample quality as it provides a detailed picture of sample heterogeneity under EM conditions while yielding initial structural insights^10^. Taken together, much can be extracted from the combined application of these techniques, but the data accuracy and resolution are limited and the associated workflows are slow, making high-throughput screening impractical. Significant challenges also arise for samples that are poor nsEM candidates or where prior structural information is unavailable. To mitigate these limitations, cryoEM-specific approaches have been developed,^4^ such as variants of the thermofluor technology^11^ and chemical crosslinking combined with density gradient centrifugation^12^.

Given that different oligomeric complexes vary in molecular mass, mass measurement could in principle be ideally suited to examine sample heterogeneity. Despite the advances in native mass spectrometry (MS) over the past decades^13–15^, the associated experimental complexity and non-native conditions have prevented native MS from becoming a widely used tool in this context. Mass photometry (MP), originally introduced as interferometric scattering mass spectrometry (iSCAMS), is a label-free approach that accurately measures molecular mass by quantifying light scattering from single biomolecules in solution^16,17^. The principle of operation of MP is remarkably similar to that of nsEM (**Fig. 1a**), where placement of a small amount (<10 μL) of solution on a substrate leads to non-specific adsorption at a solid-solute interface. In MP, in contrast to nsEM, a standard microscope cover glass replaces the carbon grid, and no stain needs to be applied.

**Figure 1.**
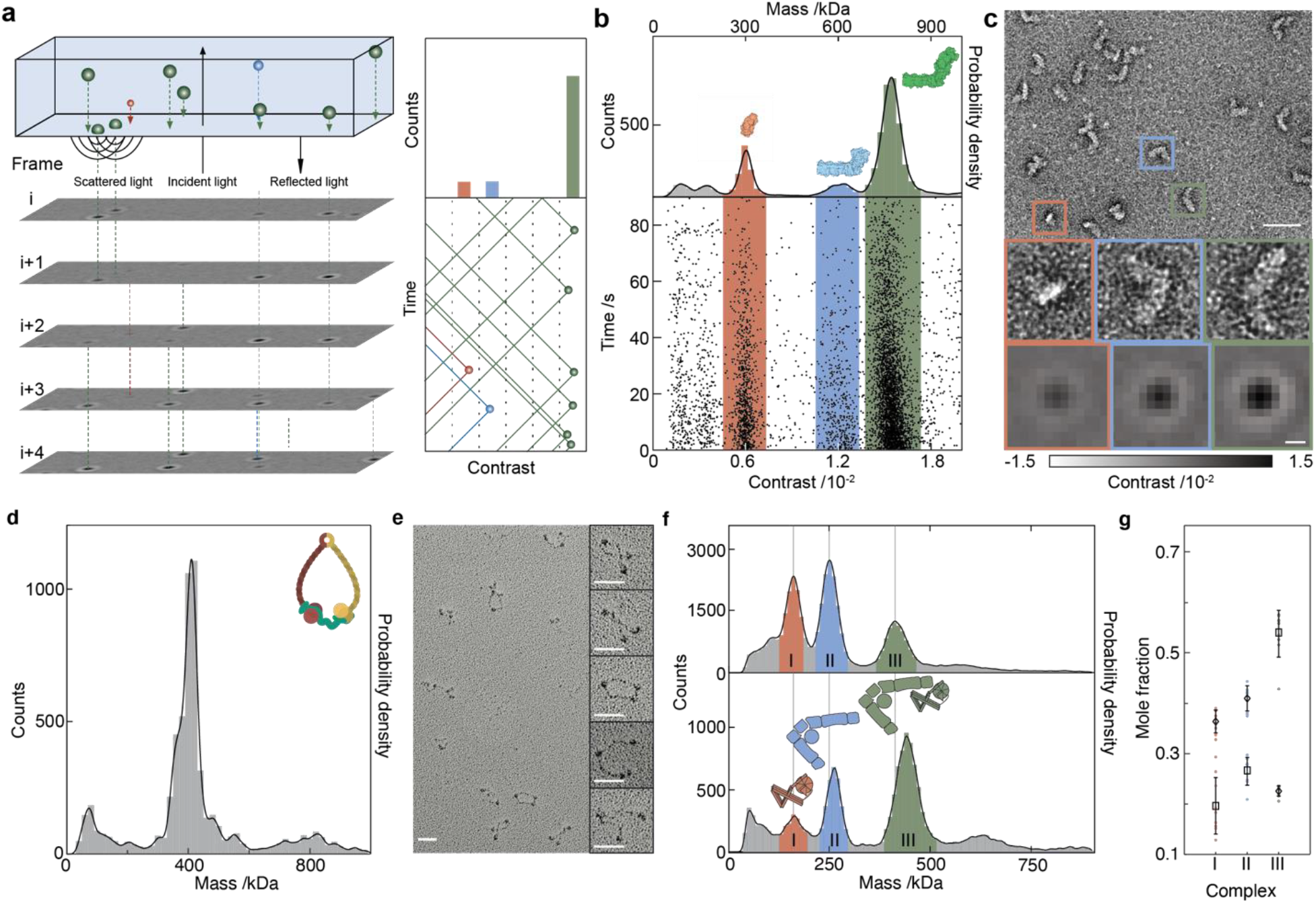
Mass photometry as a general method for characterizing biomolecular heterogeneity. **a**, Principle of operation based on interference between scattered and reflected light combined with ratiometric imaging. **b**, Scatter plot of binding events for NADH:ubiquinone oxidoreductase (respiratory complex I, 12.5 nM), and the corresponding mass histogram. **c**, Negative stain micrograph of the same sample with individual species corresponding to the peaks in **b** highlighted. MP images of species with the respective mass are shown for comparison. Scale bars: 50 nm (nsEM) and 200 nm (MP). **d**, Mass distribution for 21 nM trimeric cohesion; upper right, cartoon of trimeric cohesin. **e**, Rotary shadowing EM micrograph of trimeric cohesin shows intrinsic conformational flexibility. Scale bars: 50 nm. **f**, A mixture of two interacting NPC-I and II subcomplexes before (top) and after (bottom) chemical crosslinking with 0.1% glutaraldehyde for 5 minutes followed by quenching. **g**, Reproducibility of the crosslinking procedure shown in **f** in terms of mole fractions of the three main species before (diamonds) and after crosslinking (rectangles).

We then quantify individual binding events by illuminating the interface between the sample and cover glass and interferometrically recording reflectivity changes caused by a modification of local refractive index when an adhering biomolecule replaces water. Continuous recording of these events results in a movie of individual proteins binding to the cover glass surface (**Supplementary Movie 1**), with species appearing and disappearing in time as a consequence of the data analysis procedure^18,19^. Optimization of the image contrast^18^ then enables very accurate quantification of the reflectivity change caused by single molecule events, ultimately resulting in exceptional mass accuracy, resolution and precision^16,18^ (**Supplementary Fig. 1**).

To test the applicability of MP, we chose NADH:ubiquinone oxidoreductase (respiratory complex I) from *Escherichia coli*, a membrane-bound proton pump consisting of six soluble proteins assembled and bound to a large trans-membrane domain of seven different proteins. A scatter plot of single molecule signals arising from a recording of binding events reveals clear bands corresponding to the fully assembled, as well as partially disassembled species (**Fig. 1b**). We can convert the recorded single molecule signals to molecular mass with about 2% mass accuracy by performing a calibration routine with biomolecules of known mass (**Supplementary Fig. 2**)^16^. The resulting mass distribution shows that the majority of molecules are indeed in the fully assembled state at a complex mass of 770 kDa, in excellent agreement with previous results based on analytical ultracentrifugation^20^. The sub-complex at 600 kDa lacks the substrate acceptor module NuoEFG, while the 300 kDa species corresponds to the hydrophilic portion of the protein only^21^ (**Fig. 1b, Supplementary Fig. 3a**). A negative stain micrograph of the same sample qualitatively confirms the recorded mass and structural heterogeneity (**Fig. 1c**).

We next examined a trimeric sub-complex of cohesin, containing human SMC1, SMC3 and SCC1 fused to a C-terminal Halo-tag with a predicted molecular weight of 397 kDa. SMC1 and SMC3 contain long flexible coiled-coils, which can switch between a 50 nm extended conformation and a compacted 25 nm conformation. This means that the complex exhibits structural as well as mass heterogeneity, which would occur if the trimers were disassembled or aggregated^22,23^. The long coiled-coils and the associated structural flexibility make this complex difficult to quantify by conventional nsEM, instead requiring rotary shadowing EM. This approach demonstrates some flexibility of the coiled-coils but provides hardly any information on the integrity of the complex. MP, on the other hand, reveals a largely monodisperse sample dominated by a peak at 410 kDa, in good agreement with the expected mass, indicating that this particular sample is of excellent biochemical quality (**Fig. 1d,e; Supplementary Fig. 3b**).

We then explored whether MP could be used to evaluate efforts to improve sample purity. Such approaches – aimed at enriching the species of interest or finding appropriate stabilizing conditions – are often used when sample purity is found to be insufficient for structural characterization. These efforts, however, incur significant expense in the form of repeated staining, imaging and classification. To test whether MP could help mitigate this issue, we studied the interaction of the tetrameric Nup82-Nup145N-Nup159-Nsp1 complex (NPC-I) with a tetrameric Y-complex fragment (NPC-II) from the thermophilic fungus, *Myceliophthora thermophila*. The Y-complex forms the cytoplasmic and nucleoplasmic ring structures of the nuclear pore complex (NPC),^24,25^ a complex composed of ~500 individual proteins arranged in sub-complexes around a central eightfold rotational symmetry axis, which facilitates macromolecular transport^26^. We found that NPC-I and NPC-II bind directly and form a 1:1 complex of ~400kDa (NPC-III), which is expected given that the masses of the individual tetrameric species are 175 kDa and 259 kDa (**Fig. 1f, Supplementary Fig. 4**). After mild crosslinking with glutaraldehyde we found clear evidence for enrichment of the 1:1 complex NPC-III. In addition, we observed small increases in molecular mass for NPC-II and NPC-III. These increases were expected due to the addition of crosslinker and quencher, and the observed increases of 11 and 27 kDa were in good agreement with the number of available crosslinking sites and the subsequent quenching procedure (see Methods for full calculation). These results are highly reproducible (**Fig. 1g, Supplementary Fig. 5**), and enable us to rapidly determine how incubation time affects the resulting oligomeric distributions and, consequently, sample quality (**Supplementary Fig. 6**).

To evaluate the degree to which results obtained by MP match those from nsEM, we carried out a side-by-side comparison of nsEM and MP throughout the entire purification protocol for the Anaphase-Promoting Complex/Cyclosome (APC/C), a ubiquitin ligase essential for cell cycle progression^27–29^.

This large 1.2 MDa scaffold is composed of 19 core polypeptides and is transiently associated with numerous binding partners (**Fig. 2a, Supplementary Table 1**). Purifying homogeneous APC/C scaffolds is thus imperative for detailed biochemical and structural studies. To gain quantitative insights from nsEM, we used a map of the fully assembled APC/C that we obtained previously (**Fig. 2a**) and projected the resulting maps in-silico (**Fig. 2b**). We then used these projections as templates to assign newly generated class averages to the assembly state that they belong to.

**Figure 2.**
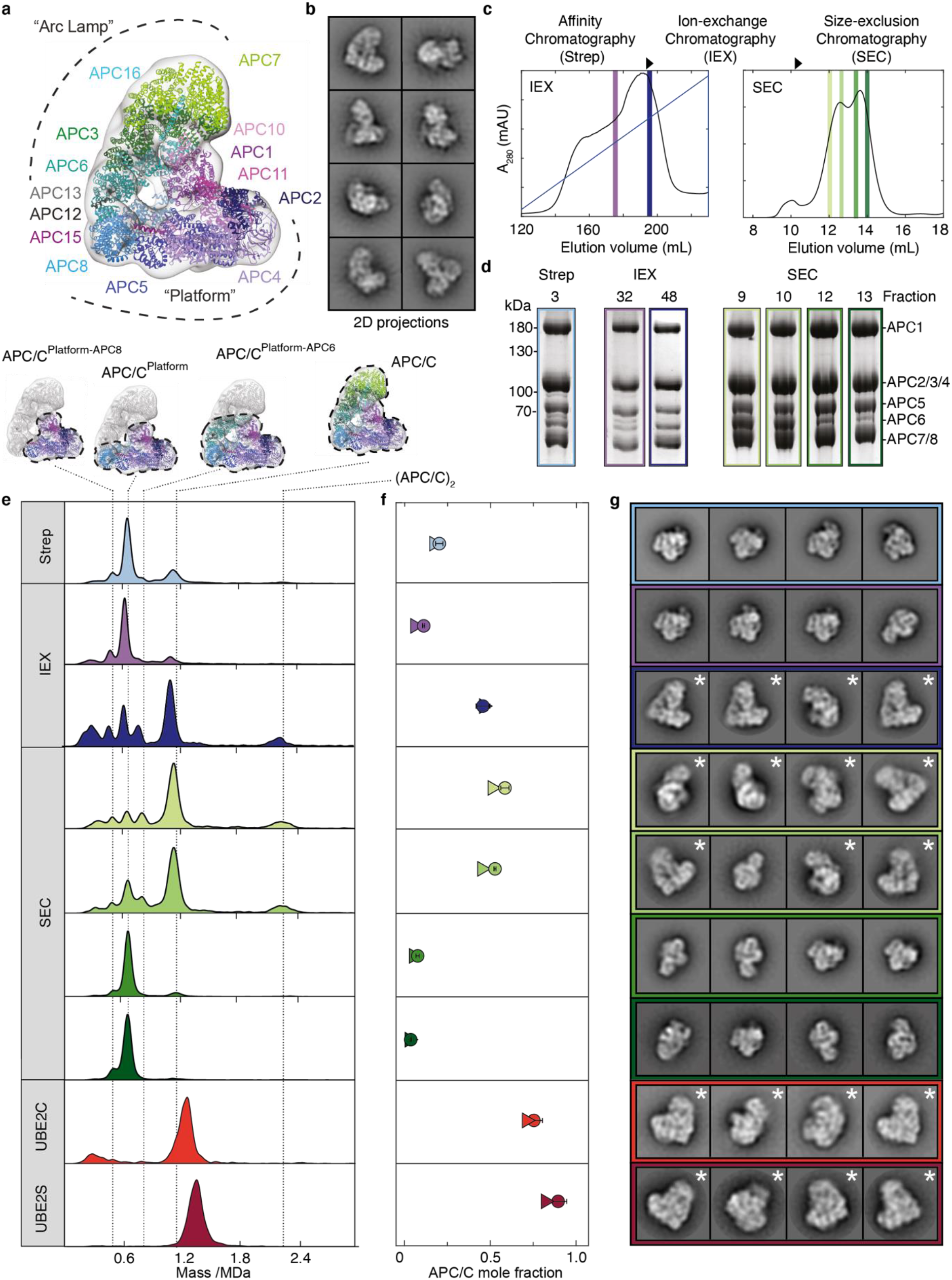
Quantitative comparison of MP with nsEM for anaphase-promoting complex/cyclosome (APC/C) purification and crosslinking. **a**, Structural cartoon including highlights of all 14 subunits. **b**, In-silico 2D projections of the fully assembled APC/C. **c**, Details of the three-step purification protocol and fractions analyzed. **d**, SDS-PAGE gels of all fractions highlighted in **c**. **e**, Corresponding mass distributions of purification step fractions and APC/C^CDH1^-UBE2C and APC/C^CDH1^-UBE2S traps indicated as UBE2C and UBE2S, respectively. **f**, Comparison of assembled fractions obtained by MP (circle) and nsEM (triangle). For the evaluation of mole fractions, we did not consider species below 400 kDa to avoid errors from buffer background for the trap samples. **g**, 2D class averages of the 4 most populated classes obtained by each step are shown. Averages representing fully assembled APC/Cs are marked by an *.

The protocol used here involves a three-step purification scheme: strep-tactin affinity, anion-exchange, and size-exclusion chromatography, with multiple fractions characterized for each of the latter two steps (**Fig. 2c**). We evaluated sample composition by SDS-PAGE (**Fig. 2d, Supplementary Figs. 7a,b,c**), mass distributions by MP (**Fig. 2e, Supplementary Figs. 7d, 8, 9**) and generated 2D classes of the same sample by nsEM in each case (**Supplementary Fig. 10)**. Strep-tactin purification yielded identical SDS-PAGE band patterns and similar mass distributions for each analyzed fraction (**Supplementary Fig. 7a,d**).

The presence of a negligible feature at 1.2 MDa suggested that only a small subset of all species was composed of fully assembled APC/C, while a sub-complex (termed “Platform”) at 660 kDa dominated (**Fig. 2e**). Quantification by nsEM conducted after strep-tactin purification confirmed the low fraction of assembled complex observed by MP (17% as measured by nsEM, 21% by MP, **Fig. 2f, Supplementary Fig. 11, Supplementary Table 2**). The top four 2D classes following strep-tactin purification, defined by selecting those with the highest number of particles out of a set of 200 2D class averages, correspondingly represent sub-complexes rather than the desired full complex (**Fig. 2g**).

Subsequent application of ion-exchange chromatography produced a mixture that still appeared heterogeneous, with the broadened elution peaks providing little information on the underlying distributions (**Fig. 2c**). SDS-PAGE of the selected fractions confirmed that the key subunits are present but revealed no discernible clues as to sample heterogeneity (**Fig. 2d, Supplementary Fig. 7b**). The MP distributions from selected fractions, however, differed dramatically, with fraction 48 demonstrating the highest contribution of fully assembled complexes following ion-exchange (45%, **Fig. 2f, Supplementary Fig. 11, Supplementary Table 2**). Accordingly, nsEM classification of fraction 48 returned fully assembled species for the top four classes, while fraction 32 consisted mainly of fragments (**Fig. 2g**). Applying this procedure to different fractions from size-exclusion chromatography demonstrated similar differences between SDS-PAGE and MP, again exhibiting quantitative agreement between MP (9: 60%, 10: 55%) and nsEM (9: 51%, 10: 45%, **Fig. 2g, Supplementary Fig. 11, Supplementary Table 2**), clearly identifying fractions 9 and 10 as optimal for structural analysis.

To evaluate the degree to which sample homogeneity can be optimized further, we explored crosslinking with two APC/C complexes, previously optimized and used for structural studies^30,31^, by MP and nsEM. These samples contained the APC/C core stably bound to a substrate (Hsl1) that is chemically linked to one of its transiently-associated cofactors (UBE2C or UBE2S), hereafter referred to as the “traps” for simplicity. These complexes were purified using tandem affinity chromatography, enriching for both a fully-assembled APC/C and the trap, and treated with GraFix^12^, glutaraldehyde crosslinking coupled with density gradient centrifugation, yielding highly purified samples (**Fig. 2e, Supplementary Fig. 8**). The agreement between MP (APC/C^CDH1^-UBE2S: 89%, APC/C^CDH1^-UBE2C: 75%, **Supplementary Fig. 11, Supplementary Table 2**) and nsEM (APC/C^CDH1^-UBE2S: 82%, APC/C^CDH1^-UBE2C: 71%) supports our previous findings^30,31^ that these purification strategies optimized sample homogeneity for cryo-EM (**Fig. 2f,g**).

The purification of protein complexes from their native source using genome editing and affinity purification strategies has become a widely-used workflow in the light of recent advances enabled by cryoEM. The intrinsic heterogeneity of the resulting complex composition, however, yields a plethora of different species in such preparations, in particular in the context of transients involving adapter proteins^32^. To explore a relevant workflow based on buffer composition rather than crosslinking, we studied the stability of the proteasome as purified from bovine heart tissue (**Fig. 3a, Supplementary Fig. 12**). The complex itself consists of two sub-complexes: The proteolytically active 20S core particle (CP) and one or two copies of the ubiquitin recognizing 19S regulatory particle (RP)^33^ (**Fig. 3b, Supplementary Fig. 13a-c, Supplementary Table 3**). Amongst others, our purification approach revealed the presence of an adapter protein called Ecm29 thought to assist the CP-RP interaction, of which we could not find any direct evidence in nsEM. The corresponding MP measurements revealed the expected features: A 2.4 MDa 30S particle (2 RP, 1 CP), a 1.5 MDa 26S particle (1RP and 1 CP),as well as the 700 kDa CP and 800 kDa RP. Additionally, we found signatures of species at both 1.7 MDa and 2.6 MDa, which we assume correspond to the 26S and 30S complexes bound to a single copy of Ecm29. Using this preparation, we screened for different buffer conditions, specifically different salt concentrations and different nucleotide states. The mass photometry distributions revealed complex disassembly with increasing salt concentration (**Fig. 3c, Supplementary Figs. 14 and 15**) as reported previously^34^, in quantitative agreement with nsEM characterization of the same samples (**Fig. 3d, Supplementary Fig. 16, Supplementary Table 4**). Interestingly, we found evidence that Ecm29 is the first to dissociate upon salt treatment, in line with previous observations that proteasome interaction proteins (PIPs) are generally salt labile^34,35^.

**Figure 3.**
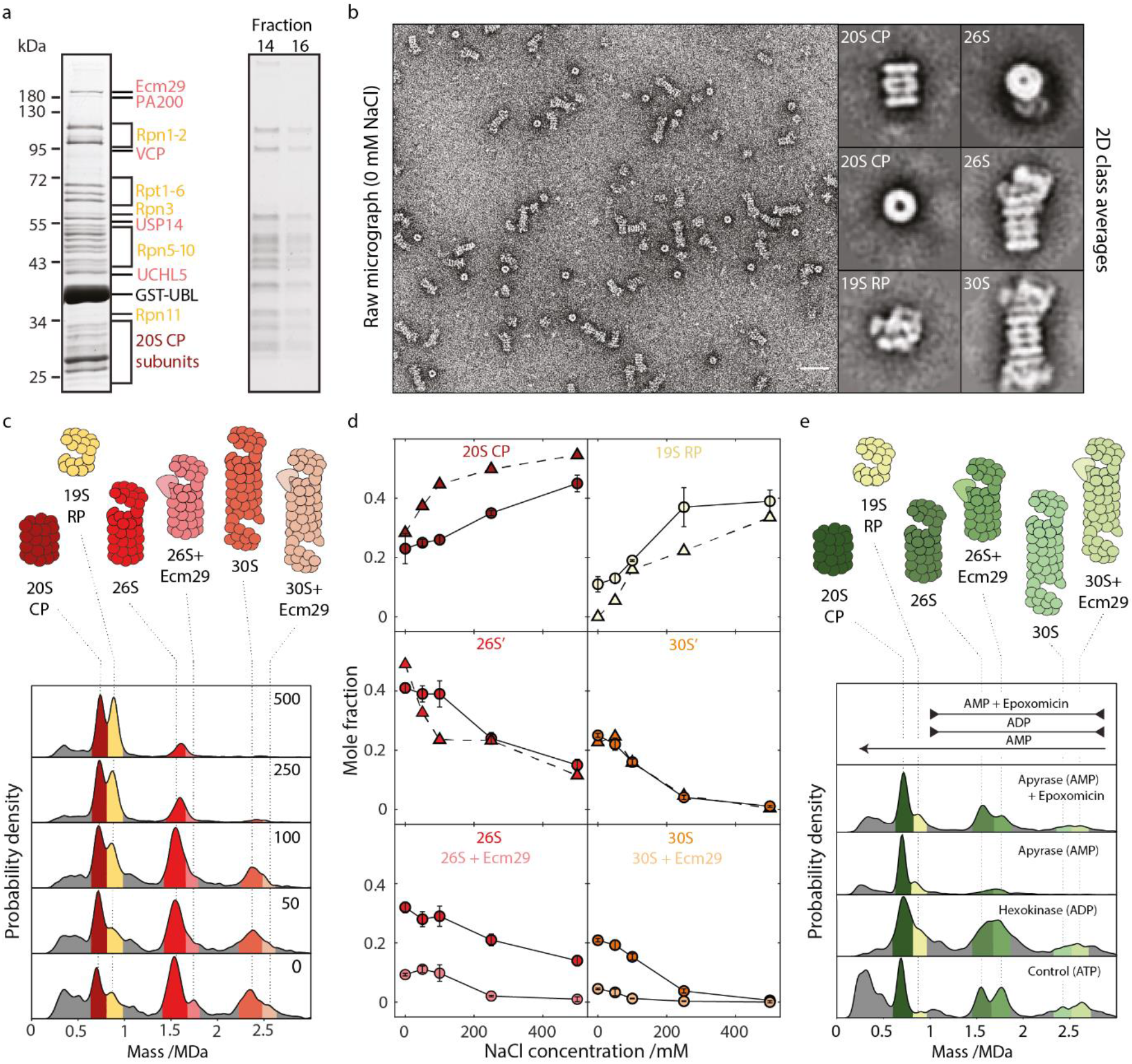
Proteasome composition, structure, stability and interactions. **a**, SDS-PAGE gels of two-step affinity purification of proteasomes – pull down with GST-Ubl (left) and separation on 10-30% sucrose gradient (right). Individual proteasome subunits are shown in dark red and yellow and PIPs in pink. **b**, Representative negative stain micrograph and 2D class averages of proteasome complexes generated by nsEM analysis. Scalebar: 50 nM. **c**, MP distributions as a function of NaCl concentration. All reactions were carried out at 4°C. **d**, Corresponding changes to the abundances of the main species comparing nsEM (triangles) with MP (circles) as well as a breakdown of the main 26S and 30S (dark) and Ecm29-bound (light) species. **e**, MP distributions as a function of different nucleotide conditions. All reactions were carried out at 37°C.

To further examine the effect of additives, we used a second preparation of bovine proteasomes and analyzed sample composition in the presence of different nucleotides. We used the enzymes apyrase, which converts ATP to AMP and hexokinase, which in the presence of glucose converts ATP to ADP. Bringing the samples to 37°C to ensure mild nucleotide exchange, resulted in considerable changes to the sub-complex distributions (**Fig. 3e, Supplementary Fig. 17**). We observed a significant increase of the 26S-bound Ecm29 fraction, which has been implicated in stabilizing the complex upon stress. The persistence of the Ecm29 complex upon apyrase treatment despite the almost complete disassembly of pure 26S agrees with the stabilizing function of Ecm29^36^. Furthermore, our data confirms that proteasome disassembly is prevented by the addition of a proteolytic inhibitor such as epoxomicin (**Fig. 3e**) most likely by a long range allosteric effect^36,37^. The observed variability in complex distributions can be confidently assigned to salt and nucleotide-induced effects, given the extreme stability of our proteasome preparations (**Supplementary Fig. 18)**.

Our results demonstrate that MP provides information on sample composition that quantitatively correlates with those obtained by nsEM (Supplementary Fig. 19), but with a number of key advantages. Measurements take place in native conditions, are extremely fast (< 1 min) and require minimal sample amounts (< pmole). The results do not rely on prior knowledge of protein structure, composition or nature of sub-complexes but instead provide direct information on which subassemblies are formed by revealing the masses of all present species. Abundances quantitatively agree with those obtained by nsEM, but are not subject to complications that can arise, such as stain artefacts,or errors in image processing including false particle picking, alignment or classification issues (**Supplementary Fig. 13d**), while revealing and quantifying species that are difficult or impossible to quantify by nsEM. Overall, MP will be of tremendous value to the cryoEM community not only by significantly improving the efficiency of structure determination, but also in applications aimed at understanding (dis)assembly processes through kinetic and reconstitution studies. More generally, the capabilities of MP will likely impact the broader life science community, by enabling accurate sample characterization for the majority of biochemical and biophysical *in vitro* studies.

## Acknowledgements

Rotary shadowing was performed by the EM Facility of the Vienna BioCenter Core Facilities GmbH (VBCF), a member of the Vienna BioCenter (VBC), Austria. All negative stain data was recorded there. Funding: A.S.S and P.K. were supported by ERC grant PHOTOMASS 819593. T.B was supported by NIH T32GM00857 & NSF DGE-1650116. G.Y. was supported by a Zvi and Ofra Meitar Magdalen Graduate Scholarship. B.A.S was supported by NIH R37GM065930, NIH P30CA021765, American Lebanese Syrian Associated Charities, and Howard Hughes Medical Institute HHMI. J.S. and T.F. were supported by the Deutsche Forschungsgemeinschaft grants -278002225/RTG 2202 and FR 1140/11-2. P.V.D. and T.U.S were supported by NIH R01GM77537. B.B. was supported by long term EMBO and HFSP fellowships. Research in the laboratory of J-MP is supported by Boehringer Ingelheim, the Austrian Research Promotion Agency (Headquarter grant FFG-852936) and the ERC under the European Union’s Horizon 2020 research and innovation programme GA No 693949, and by HFSP grant RGP0057/2018. N.G.B was supported by NIH R35GM128855, NIH P30AI050410, and UCRF.

## Contributions

ASS, KB, TB contributed equally and will be putting their name first on the citation in their CVs. Concept: NGB, DH and PK. Methodology: ASS, KB, TB, GY, NGB, DH, PK. Formal analysis: ASS, KB, TB, GY. Investigation: ASS, KB, TB, GY. Resources: TB, RTV, BAS, NGB, KB, DH, JS, TF, PVD, TUS, BB, JMP. Writing of original draft: ASS, JLPB, NGB, DH, PK. Revision and editing: TB, NGB, KB, DH, JS, TF, BB, JMP, PVD, TUS, ASS, GY, WBS, JLPB, PK. Visualization: ASS, KB, TB, WBS, JLPB, PK

Supervision: NGB, DH, PK

## Conflict of interest

P.K, J.L.P.B, G.Y and W.B.S are academic founders and consultants to Refeyn. All other authors declare no conflict of interest.

## Methods

### Mass Photometry

Microscope coverslips (No. 1.5, 24×50 and 24×24 mm^2^, VMR) were cleaned by sequential sonication in 50% isopropanol (HPLC grade)/Milli-Q H_2_O, and Milli-Q H_2_O (5 minutes each), followed by drying with a clean nitrogen stream. Clean coverslips were assembled into flow chambers using double-sided-sticky tape (3M) as described by Young et al^16^. Fresh aluminium foil was folded around an A4 size cutting board. Individual 24×24 coverslips were taped using two strips of double-sided tape and cut from the foil using a scalpel blade. Each excised 24×24 coverslip was joined, tape side down, in the center of a 24×50 coverslip and stored prior to use.

Immediately prior to mass photometry measurements, protein stocks were diluted directly in stock buffer (unless stated otherwise). Typical working concentrations of protein complexes were 5-25 nM, depending on the dissociation characteristics of individual assemblies. Each protein was measured in new flow-chambers (i.e. each flow-chamber was used once). To find focus, fresh buffer was first flowed into the chamber, the focal position was identified and secured in place with an autofocus system based on total internal reflection for the entire measurement. For each acquisition, 15 μL of diluted protein was introduced into the flow-chamber and, following autofocus stabilization, movies of either 60 or 90 s duration recorded. Each sample was measured at least 3 times independently (*n* ≥ 3).

All measurements were performed using similar mass photometry instruments. Most data was acquired using a ONE^MP^ mass photometer (Refeyn LTD, Oxford, UK) except for nuclear pore complex (NPC) crosslinking experiments which were performed on a home-built mass photometer operating at the same wavelength as the commercial instrument. Data acquisition was performed using either AcquireMP (Refeyn LTD, v1.1.3 – proteasome measurements and v1.2.1 for all other measurements) or custom software written in Labview (for NPC crosslinking^16^). Mass photometry movies were recorded at 1 kHz, with exposure times varying between 0.6-0.9 ms, adjusted to maximize camera counts while avoiding saturation. Images were time averaged 5-fold and pixel binned 4×4, before saving. The time and pixel binning resulted in an effective pixel size of 84.4 nm and effective frame rate of 200 Hz.

| **Protein** | **Number of**<br>**binned**<br>**frames, $nf$** | **Threshold**<br>**1** |
| --- | --- | --- |
| Respiratory complex I | 5 | 0.5 |
| Trimeric cohesin | 10 | 0.1 |
| NPC | 5 | 0.8 |
| APC/C – affinity chromatography | 3 | 2 |
| APC/C – ion exchange chromatography | 5 | 3 |
| APC/C – size-exclusion chromatography | 3 | 2 |
| APC/C – crosslinked | 3 | 2 |
| Proteasome | 5 | 3.5 |

DiscoverMP analysis parameters for all protein complexes. The values for threshold 2 (= 0.25) and median filter kernel (= 15) remained constant for all protein samples.

#### Image processing

All MP images were processed and analyzed using DiscoverMP (v1.2.3). In short, the procedure included three main steps: 1) background removal, 2) identification of landing particles and 3) particle fitting to extract maximum contrast. *Background removal* – static scattering background from the glass-water interface was removed by calculating the ratiometric images, *R*, as *R_m_* = *N_m_*/*N*_*m*-1_ – 1, where 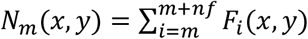 is the sum of each pixel in consecutive images (*F*), with *nf* defining how many frames to sum^16^. In this way, images in the field of view are preserved, while eliminating any background. This procedure is applied to all possible frames, creating a ratiometric movie (**Supplementary Movie 1**), where the binding of particles to the glass-water interface is clearly visible. *Identification of landing particles* – a landing particle generates a step-wise increase in the glass reflectivity which results in an increase in scattering signal, followed by an amplitude decrease ratiometric movie^16^. This distinct signature of step-wise behavior is used to identify particles, using two fitting parameters; threshold 1 (related to the particle contrast relative to the background noise) and threshold 2 (related to the radial symmetry or the particle signature). *Particle fitting* – identified particles were fit using a model point spread function (PSF) in order to extract the contrast. **Supplementary Fig. 1** shows the experimental and fitted PSF with the corresponding residuals, emphasizing the accuracy of the fitting procedure, both for particles with large (top panel) and small (bottom panel) signal-to noise ratio.

#### Calibration procedure

Contrast-to-mass (C2M) calibration was performed daily and for each buffer solution separately, since the C2M conversion may change slightly as a result of buffer content. The calibration protocol included measurement of two protein oligomer solutions, one with masses of 66, 132 and 198 kDa and a second with masses of 90, 180, 360 and 540 kDa. Each MP calibration experiment was analyzed using DiscoverMP. The mean peak contrast was determined in the software using Gaussian fitting. The mean contrast values from both calibration protein solutions were then plotted (**Supplementary Fig. 2**) and fitted to a line, *y* = *bx*, with *y* – contrast, *x* – mass and *b* – C2M calibration factor.

#### Extraction of mole fractions

The output from the analysis of each individual movie resulted in a list of individual particle contrasts, which were converted to mass using the corresponding C2M calibration. From these data, kernel density estimates (KDE) were generated for each sample using a Gaussian kernel with a fixed bandwidth using MATLAB (R2017b), which helped eliminate variations due to total particle counts between experiments. Bandwidth values varied between the different proteins, and were determined by experimental noise, where noisier data required larger bandwidths. Bandwidth values were 15 (Cohesin and Complex I), 20 (APC/C and NPC) and 30 (proteasome) kDa. To estimate the mole fractions of the different species, the KDEs were then fitted to a sum of several Gaussian functions. The number of Gaussians chosen as well as the respective center of mass values were identified based on the apparent number of peaks in the KDE plots. This included the presence of peak shoulders and the existence of (sub-) complexes in a given sample. We fitted the Gaussian sum using MATLAB curve fitting tools. The relative amount of each species was calculated as the area of each Gaussian (i.e., 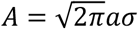, with *A* – area, *a* – amplitude and *σ* – standard deviation of the fitted Gaussian). Only relevant species were chosen and the mole fractions were calculated by renormalizing the area from the area sum of all relevant species.

All NPC crosslinking measurements were fitted to three Gaussians at 160, 260 and 410 kDa for experiments without crosslinking, and at 160, 270 and 440 kDa for the crosslinking ones (See **Supplementary Fig. 5** for a typical example). All APC/C purification step analyses were fitted to four Gaussians at 490, 640, 800, 1175 kDa corresponding to the main (sub-) complex species we observed (See **Supplementary Fig. 9** for a typical example). All APC/C crosslinked samples were fitted to a single Gaussian. All Proteasome salt addition analyses were fitted to seven Gaussians at 720, 890, 1050, 1550, 1750, 2350, 2550 kDa (See **Supplementary Fig. 15** for a typical example). The mass difference in the 26S peaks (1550/1750 kDa) and 30S peaks (2350/2550 kDa) arises from proteasome binding partners that co-purify with proteasomes in a sub-stochiometric amounts. Since negative staining analysis cannot easily distinguish between the different species of 26S and 30S with/without co-factors bound, we summed up the respective Gaussians to represent the amount of 26S and 30S species in the solution that are classified together in negative stain analysis. In all cases described above, all repeats of measurements were fitted separately, subsequently estimating the mole fraction values, followed by a calculation of the mean and standard deviation (estimated measurement error).

#### Correction for surface-solution concentrations discrepancies

In mass photometry proteins bind to a surface, and thereby decrease the overall concentration of the protein in solution. Young et al^16^ showed that the main factor affecting the binding rate of different protein (sub-)complexes to the surface is their diffusion^16^, characterized by an exponential decay in time with a rate constant roughly proportional to (molecular weight)^−1/3^. As described above, every MP measurement starts following autofocus stabilization, and therefore proteins that bind within this ‘deadtime’ (approximately 10-15 seconds) are not recorded. This is in contrast to negative staining where particles information is ‘recorded’ from the moment the droplet is placed on the surface. This discrepancy suggests that MP measurements underestimate the abundance of smaller proteins, as compared to negative staining. Since smaller proteins diffuse faster to the surface, they are likely to be more depleted during the ‘dead-time’, and as a result a relative shift towards higher mass distributions may be observed. This potential shift can be easily accounted for by applying a *diffusion correction* to the counted numbers of each protein species depending on their binding rate^16^. This correction is based on the comparison between the integration over the exponential decay in binding events from sample addition (at time zero, *t* = 0) until the end of the measurement time (*t* = *t_f_*), and the integration over the real experimental time, which starts at *t* = *t*_0_ and ends at *t* = *t_f_*:

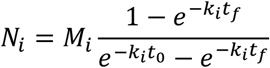

where *N_i_* is the number of particles of species *i* counted over the full integral (*t* = 0 – *t_f_*), *M_i_* is the experimentally measured number of particles, and *k_i_* is the binding rate constant for species *i* and is proportional to a given molecular weight (MW), *k_i_* ≅ *α* · (*MW_i_*)^−1/3^. The proportionality prefactor, *α*, was calculated from the landing rate of all particles in a movie and their weighted average mass, and was very similar across all movies (*α* = 0.2 and 0.3 s^−1^ · kDa^1/3^ for proteasome and APC/C experiments, respectively). The correction was applied to all APC/C purification step measurements (**Supplementary Fig. 11**) and proteasome salt addition experiments (**Supplementary Fig. 16**), and compared to the negative staining distributions. Crosslinked APC/C mole fractions were not diffusion-corrected as their KDEs exhibit only one distinct Gaussian peak. As expected, raw, uncorrected, distributions underestimated the abundance of smaller species compared to nsEM (**Supplementary Fig. 11** – APC/C, **Supplementary Fig. 16** – proteasome), while the corrected distributions exhibit excellent agreement. Supplementary Tables 2, 4 summarize the raw and corrected mole fraction values (**Supplementary Table 2** – APC/C, **Supplementary Table 4** – proteasome).

#### Crosslinking NPC-III protein

For the crosslinking of NPC-III, the protein was incubated at a concentration of 0.18 mg/mL (434 nM of complex) with 0.1% glutaraldehyde for 5 minutes on ice in a total volume of 9 μL, before being quenched with 1 μL of quenching buffer (crosslinking protocol was inspired from^38^). Samples taken from this final, quenched, solution were diluted 10-fold in buffer immediately before mass photometry measurements. From each reaction volume, three independent measurements were taken, with the premeasurement dilution performed separately for each one. Three independent reactions were carried out, resulting in a total of 9 measurements of the crosslinked species. For the measurements of the complex without crosslinking, the complex was diluted 10-fold from the stock concentration of 0.2 mg/mL (482 nM of complex), again immediately before measurement. This procedure was repeated for 9 independent measurements of the complex without crosslinker. Crosslinking experiments were also performed as a function of incubation time. For this, the procedure described above was repeated with incubation times of 5, 10, 15 and 20 mins (**Supplementary Fig. 6**). For each time point, 2 measurements were taken, and 2 measurements of the complex straight after dilution without crosslinking were also taken. The buffer used throughout was 10 mM HEPES, pH 7.5, 150 mM NaCl, 1 mM DTT and 0.1 mM EDTA. The quenching buffer contained 8 mM aspartate (Asp) and 2 mM lysine (Lys).

#### Mass shift due to crosslinking of NPC-III protein

There are 160 lysine residues (lys) in NPC-III. Assuming all those residues bind a glutaraldehyde molecule (100 Da), the full complex mass should increase by 16 kDa. The quenching buffer includes 0.8mM Asp (133 Da) and 0.2mM Lys (146 Da), which can each quench glutaraldehyde, adding another ~22 kDa per NPC-III assembly. In total, the crosslinking procedure could add up to ~38 kDa per NPC-III assembly. The observed mass shift in our experiments was 27 kDa (**Fig. 1f**), within the expected range.

### Negative Staining

*All negative staining grids were prepared with the exact protein samples and in the same concentrations used for MP measurements.*

#### Negative stain sample preparation, data collection and image processing – APC/C and Proteasome

The samples were stained with 2 % (w/v) uranyl acetate. Carbon-coated grids were glow-discharged using EM ACE600 sputter coater (*Leica*) for 30 seconds at ~20 mA. 4 μL of the sample was applied on the glow-discharged grid and incubated for several seconds. The excess liquid was blotted off using filter paper. The grid was washed three times with a water droplet. 4 μL of 2% uranyl acetate was applied on the grid with adsorbed sample and incubated for 1 minute. The excess stain was blotted off using a filter paper. The grids were air-dried before micrograph recording. Images were recorded on FEI Tecnai G^2^ 20 (*FEI*) transmission electron microscope at a magnification of 62k× (1.85 Å/pixel). Particle picking was done using CrYOLO^39^ after which the particles were transferred to cowEyes(https://www.cow-em.de/) for subsequent rounds of 2D classifications. After final 2D classification, clean 2D class averages were extracted and representative 2D classes were visualized using RELION v3.0^40^.

#### Rotary shadowing imaging – Cohesin

Cohesin trimer was first diluted to a concentration of approximately 0.1 mg/mL in 50 mM sodium phosphate buffer pH 7.6 (including 150 mM NaCl, 5% glycerol and 0.5mM TCEP) and subsequently diluted 1:1 in spraying buffer, containing 200 mM ammonium acetate and 60% (v/v) glycerol, pH adjusted to 7.6. After dilution, the samples were sprayed onto freshly cleaved mica chips (Agar Scientific, UK) and immediately transferred into a BAL-TEC MED020 high vacuum evaporator (BAL-TEC, Liechtenstein) equipped with electron guns. While rotating, samples were coated with 0.7 nm Platinum (BALTIC, Germany) at an angle of 4-5°, followed by 7 nm Carbon (Balzers, Liechtenstein) at 90°. The obtained replicas were floated off from the mica chips, picked up on 400 mesh Cu/Pd grids (Agar Scientific, UK), and inspected in an FEI Morgagni 268D TEM (FEI, The Netherlands) operated at 80kV. Images were acquired using an 11 megapixel Morada CCD camera (Olympus-SIS, Germany).

### Protein production, purification and measurement condition

#### Respiratory complex I

Respiratory complex I from *Escherichia coli* was prepared as described previously^41^ with slight modifications. After affinity chromatography on a Probond Ni^2+^-IDA column (35 mL), the complex was subjected to a Superose 6 (24 mL) size exclusion column. Peak fractions were concentrated to 30 μM (Amicon Ultra-15, 100 kDa MWCO) and stored in 50 mM MES/NaOH, 50 mM NaCl, 5 mM MgCl_2_ and 0.005% MNG-3 (pH 6.0) at −80°C. MP and nsEM measurements of complex I were performed at 12.5 nM, with a drop-down dilution with its storing buffer without MNG-3, and within 15 mins of dilution, to avoid any protein aggregation due to lower MNG-3 concentration.

#### Cohesin (trimers)

SF9 insect cells were transfected for 48 hrs with bacmids harboring homo sapiens (hs) SMC1, hs SMC3-FLAG and hs RAD21-Halo-His14 in the pBig1a expression vector^42^. Cells were collected by centrifugation, washed with 1xPBS and snap frozen in liquid nitrogen. Cells were lysed by douncing 25 times in Lysis Buffer (50 mM NaPO_4_, pH 7.6, 500 mM NaCl, 5% glycerol, 0.05% Tween, 10 mM Imidazole) supplemented with EDTA-free protease inhibitor cocktail (Roche), 1 mM phenyl-methyl-sulfonyl-fluoride (PMSF), 1 mM benzamidine, and 3 mM beta-mercapto-ethanol (bME). Insoluble material was removed by centrifugation (18.5 krpm, LYNX, 35 min, 4°C) and the supernatant applied for 3 h at 4°C to 5 mL of Toyopearl AF-Chelate-650M resin (Tosoh) precharged with Ni^2+^-ions. Beads were collected by low speed centrifugation, washed in batch two times with 10 column volumes (CVs) of Lysis buffer supplemented with 5 mM imidazole, collected in a glass column in 5 CVs of Lysis buffer plus 5 mM imidazole and eluted with 5 CVs Elution buffer (50 mM NaPO_4_, pH 7.6, 150 mM NaCl, 5% glycerol, 300 mM Imidazole). The eluate was incubated for 3 h at 4°C with 5 mL of FLAG M2 Agarose beads (Sigma), collected in a glass column, washed with 2×5 CVs of FLAG-Buffer (50 mM NaPO_4_, pH 7.6, 100 mM NaCl, 5% glycerol, 50 mM Imidazole) and eluted with 5×1 CV of FLAGbuffer supplemented with 0.25 mg/mL 3×FLAG peptide. The eluate was bound to 75 μL of POROS HS resin (Thermo) for 30 min at 4°C. The beads were collected in a disposable plastic column, and bound cohesin eluted with 3×150 μL High Salt Buffer (50 mM NaPO_4_, pH 7.6, 750 mM NaCl, 5% glycerol, 50 mM Imidazole, 0.5mM TCEP). The eluate was dialyzed over night against Dialysis Buffer (50 mM NaPO_4_, pH 7.6, 100 mM NaCl, 5% glycerol, 0.5 mM TCEP), aliquoted, snap-frozen in liquid nitrogen and stored at −80°C. Protein concentrations were determined by the Bradford Assay, assuming a molecular weight of 397 kDa. MP measurements of Cohesin trimers were performed at 21 nM, diluted from stock solution using its storing buffer.

#### Nuclear Pore Complex

##### Construct generation

###### NPC-I

Nup82_1–854_, Nup159_1072–1447_ and Nsp1_507–718_ were cloned from GeneArt Strings (ThermoFisher Scientific) into modified pET-Duet plasmids containing three expression cassettes. The first cassette contained Nup159_1072–1447_ fused to an N-terminal TEV protease cleavable His6-Avi-MBP tag. Nup82_1–854_ and Nsp1_507–718_ were cloned into the second and third cassette, respectively, without any modification. Nup145N_868–1004_ was cloned from *M. thermophila* into a pET plasmid introducing an N-terminally fused human rhinovirus 3C (3C)–cleavable His_10_-Arg_8_-SUMO tag. The tetrameric complex, Nup82-Nup159cc-Nsp1cc-Nup145N_APD_ is referred to as NPC-I.

###### NPC-II

Nup120_952–1241_, Nup145C_1005–1791_, Nup85_257–1181_ and full-length Sec13 were cloned from *M. thermophila*. To increase stability, Sec13 was spliced between Nup145C_1005–1232_ and Nup145C_1233–1791_ to generate a structure-based fusion protein. The Nup145C_1005–1232_-Sec13-Nup145C_1233–1791_ protein construct was N-terminally tethered to a 3C-cleavable SUMO tag. Nup85_257–1181_ was N-terminally fused with a 3C–cleavable His_10_-Arg_8_-SUMO. Nup120_952–1241_ was C-terminally fused with a His_10_ tag. All constructs were cloned into pET-derived plasmids. The tetrameric complex Nup120-Nup145C-Sec13-Nup85 is referred to as NPC-II for simplicity.

##### Protein production and purification

*Escherichia coli* LOBSTR-RIL(DE3) (Kerafast)^43^ cells were co-transformed with vectors, and protein production was induced with 0.2 mM IPTG at 18°C for 12–14 h.

##### Production of NPC-III, the octameric Nup82-Nup159cc-Nsp1cc-Nup145N_APD_-Nup120-Nup145C-Secl3-Nup85

NPC-II was purified as in^44^. Cells expressing trimeric Nup82_1–854_-Nsp1_507–718_-Nup159_1072–1447_ were collected by centrifugation at 6,000*g*, resuspended in lysis buffer (50 mM potassium phosphate, pH 8.0, 500 mM NaCl, 3 mM β-mercaptoethanol (βME) and 1 mM PMSF) and lysed with an LM20 microfluidizer (Microfluidics). The lysate was cleared by centrifugation at 12,500*g* for 25 min. The soluble fraction was incubated with amylose resin (NEB) for 30 min at 4°C. After washing of the beads with lysis buffer, the protein was eluted (10 mM maltose, pH 8.0, 150 mM NaCl and 3 mM βME). The Nup82_1–854_-Nsp1_507–718_-Nup159_1072–1447_ amylose eluate was incubated with TEV and dialyzed overnight at 4°C (10 mM Tris-HCl, pH 8.0, 150 mM NaCl, 0.1 mM EDTA and 1 mM DTT). Purified Nup145NAPD was incubated with trimeric Nup82_1–854_-Nsp1_507–718_-Nup159_1072–1447_ and the assembled tetrameric NPC-I complex separated by size-exclusion chromatography on Superdex S200 (GE Healthcare) 20 mM HEPES-KOH, pH 7.4, 0.1 mM EDTA and 1 mM DTT). NPC-I and NPC-II were mixed and NPC-III isolated by size-exclusion chromatography on Superdex S200 (20 mM HEPES-KOH, pH 7.4, 0.1 mM EDTA and 1 mM DTT).

#### APC/C purification and sample preparation

Recombinant APC/C was expressed in High Five insect cells (Thermo Scientific) as described in^30,42^. Briefly, APC/C was expressed with a Twin-Strep(II)-tag on the C-terminus of APC4 and purified using Strep-Tactin Sepharose (IBA Life Sciences) affinity chromatography followed by ion exchange chromatography and SEC. Final buffer conditions were 20 mM Hepes pH 8, 200 mM NaCl, 1 mM DTT. Crosslinked APC/C EM samples were prepared as described in^30,31^.

All APC/C samples were diluted to working concentration using APC/C buffer (20 mM Hepes pH 8, 200 mM NaCl), which varied between samples to optimize background noise and particles counts. The concentrations were Strep (3, 4, 5) 12 nM, IEX (26, 32, 38, 48) 5 nM, SEC (9, 10, 12, 13) 15 nM. APC/C^CDH1^-UBE2C and APC/C^CHD1^-UBE2S traps were measured directly after buffer exchange using a Zeba spin column (Pierce) to remove glycerol.

#### Proteasome

Proteasome complexes were purified from bovine heart extract by the protocol adapted from^45^. Briefly, bovine heart was homogenized in purification buffer (25 mM BisTris, pH 6.5, 50 mM KCl, 5 mM MgCl_2_, 10% glycerol, 4 mM ATP and 1 mM DTT) and cleared in Optima XE-90 Ultracentrifuge, Ti45 rotor (*Beckman Coulter*) for 1 h at 100000g at 4°C. The final extract was prepared by two-step protein precipitation with 4% and 20% PolyEthyleneGlycol8000 (PEG8000). Precipitated proteins were dissolved in purification buffer. Proteasomes were affinity purified with the bait protein GST-Ubl and magnetic beads (MagneGST™ Glutathione Particles, *Promega)* and eluted with purification buffer containing 25 mM reduced L-glutathione. Proteasome samples were concentrated and applied on a 10–30% sucrose gradients (purification buffer containing 10 or 30% (w/v) sucrose). Gradients were run in Optima XE-90 Ultracentrifuge, SW60Ti rotor (*Beckman Coulter*) for 16 hours at 100000g and 4°C. Gradients were manually fractionated into 200 μL fractions and protein concentrations were determined by the Bradford assay (Protein Assay Dye Reagent Concentrate 5×, *Bio-Rad*).

#### Sample preparation under different salt concentrations/nucleotide conditions

Prior to mass photometry measurements, proteasome samples (1 μM) were buffer exchanged (Zeba Micro Spin Desalting Columns, *Thermo Fisher Scientific*) into 25 mM BisTris, 50 mM KCl, 5 mM MgCl_2_, 20% glycerol, 4 mM ATP and 1 mM DTT and diluted 2x. For measuring proteasome stability in the presence of NaCl, the proteasome sample was first diluted to 50 nM and then NaCl was added to a final concentration of 50, 100, 250 and 500 mM, and samples were incubated on ice for 2 hours. For measuring proteasome stability under different nucleotide conditions either apyrase (100 mU), which hydrolyzes ATP and ADP to AMP, hexokinase (100 mU) and D-glucose (20 mM), which hydrolyses ATP to ADP in a reaction that results in generation of glucose-6-phosphate, apyrase (100 mU) and proteasome inhibitor epoxomicin (50 μM), which stabilized proteasome complex were added to 2x diluted proteasomes and incubated for 2 hours at 37°C. Before measurements, proteasome samples were diluted to 50 nM.

## References

1. Pieters, B. J. G. E., Van Eldijk, M. B., Nolte, R. J. M. & Mecinović, J. Natural supramolecular protein assemblies. Chem. Soc. Rev. 45, 24–39 2016.

2. Peng, H. P., Lee, K. H., Jian, J. W. & Yang, A. S. Origins of specificity and affinity in antibody-protein interactions. Proc. Natl. Acad. Sci. U. S. A. 111, 2014.

3. Robinson, C. V, Sali, A. & Baumeister, W. The molecular sociology of the cell. Nature 450, 973–982 2007.

4. Passmore, L. A. & Russo, C. J. Specimen Preparation for High-Resolution Cryo-EM. Methods in Enzymology 579, (Elsevier Inc., 2016).

5. Vinothkumar, K. R. & Henderson, R. Single particle electron cryomicroscopy: trends, issues and future perspective. Q. Rev. Biophys. 49, 2016.

6. Glaeser, R. M. How Good Can Single-Particle Cryo-EM Become? What Remains Before It Approaches Its Physical Limits? Annu. Rev. Biophys. 48, 45–61 2019.

7. Campos-Acevedo, A. A., Díaz-Vilchis, A., Sotelo-Mundo, R. R. & Rudiño-Piñera, E. First attempts to crystallize a non-homogeneous sample of thioredoxin from Litopenaeus vannamei: What to do when you have diffraction data of a protein that is not the target? Biochem. Biophys. Reports 8, 284–289 2016.

8. Oliveira, C. & Domingues, L. Guidelines to reach high-quality purified recombinant proteins. Appl. Microbiol. Biotechnol. 102, 81–92 2018.

9. Gast, K. Dynamic and Static Light Scattering. in Instrumental Analysis of Intrinsically Disordered Proteins: Assessing Structure and Conformation 477–524 2010. doi:10.1002/9780470602614.ch17

10. Ohi, M., Li, Y., Cheng, Y. & Walz, T. Negative Staining and Image Classification – Powerful Tools in Modern Electron Microscopy. Biol. Proced. Online 6, 23–34 2004.

11. Chari, A. et al. ProteoPlex: Stability optimization of macromolecular complexes by sparse-matrix screening of chemical space. Nat. Methods 12, 2015.

12. Kastner, B. et al. GraFix: sample preparation for single-particle electron cryomicroscopy. Nat. Methods 5, 53–55 2007.

13. Chorev, D. S. et al. Protein assemblies ejected directly from native membranes yield complexes for mass spectrometry. Science 362, 829–834 2018.

14. Donnelly, D. P. et al. Best practices and benchmarks for intact protein analysis for top-down mass spectrometry. Nat. Methods 16, 587–594 2019.

15. Ishii, K., Zhou, M. & Uchiyama, S. Native mass spectrometry for understanding dynamic protein complex. Biochim. Biophys. Acta – Gen. Subj. 1862, 275–286 2018.

16. Young, G. et al. Quantitative mass imaging of single biological macromolecules. Science 360, 423–427 2018.

17. Young, G. & Kukura, P. Interferometric Scattering Microscopy. Annu. Rev. Phys. Chem. 70, 301–322 2019.

18. Cole, D., Young, G., Weigel, A., Sebesta, A. & Kukura, P. Label-Free Single-Molecule Imaging with Numerical-Aperture-Shaped Interferometric Scattering Microscopy. ACS Photonics 4, 211–216 2017.

19. Piliarik, M. & Sandoghdar, V. Direct optical sensing of single unlabelled proteins and superresolution imaging of their binding sites. Nat. Commun. 5, 1–8 2014.

20. Böttcher, B., Scheide, D., Hesterberg, M., Nagel-Steger, L. & Friedrich, T. A novel, enzymatically active conformation of the Escherichia coli NADH:ubiquinone oxidoreductase (complex I). J. Biol. Chem. 277, 17970–17977 2002.

21. Leif, H., Sled, V. D., Ohnishi, T., Weiss, H. & Friedrich, T. Isolation and Characterization of the Proton-translocating NADH:ubiquinone Oxidoreductase from Escherichia coli. Eur. J. Biochem. 230, 538–548 1995.

22. Hons, M. T. et al. Topology and structure of an engineered human cohesin complex bound to Pds5B. Nat. Commun. 7, 1–11 2016.

23. Bürmann, F. et al. A folded conformation of MukBEF and cohesin. Nat. Struct. Mol. Biol. 26, 227–236 2019.

24. Von Appen, A. et al. In situ structural analysis of the human nuclear pore complex. Nature 526, 140–143 2015.

25. Lin, D. H. et al. Architecture of the symmetric core of the nuclear pore. Science 352, 2016.

26. Knockenhauer, K. E. & Schwartz, T. U. The Nuclear Pore Complex as a Flexible and Dynamic Gate. Cell 164, 1162–1171 2016.

27. Watson, E. R., Brown, N. G., Peters, J. M., Stark, H. & Schulman, B. A. Posing the APC/C E3 Ubiquitin Ligase to Orchestrate Cell Division. Trends Cell Biol. 29, 117–134 2019.

28. Yamano, H. APC/C: current understanding and future perspectives. F1000 Research 8, 1–14 2019.

29. Alfieri, C., Zhang, S. & Barford, D. Visualizing the complex functions and mechanisms of the anaphase promoting complex/cyclosome (APC/C). Open Biol. 7, 2017.

30. Brown, N. G. et al. Dual RING E3 architectures regulate multiubiquitination and ubiquitin chain elongation by APC/C. Cell 165, 1440–1453 2016.

31. Brown, N. G. et al. RING E3 mechanism for ubiquitin ligation to a disordered substrate visualized for human anaphase-promoting complex. Proc. Natl. Acad. Sci. U. S. A. 112, 5272–5279 2015.

32. Tai, H.-C., Besche, H. C., Goldberg, A. L. & Schuman, E. M. Characterization of the brain 26S proteasome and its interacting proteins. Front. Mol. Neurosci. 3, 1–19 2010.

33. Peters, J.-M., Harris, J. R. & Kleinschmidt, J. A. Ultrastructure of the ~26S complex containing the ~20S cylinder particle (multicatalytic proteinase/proteasome). Eur. J. Biochem. 56, 422–432 1991.

34. Leggett, D. S. et al. Multiple Associated Proteins Regulate Proteasome Structure and Function. Mol. Cell 10, 495–507 2002.

35. Besche, H. C., Haas, W., Gygi, S. P. & Goldberg, A. L. Isolation of mammalian 26S proteasomes and p97/VCP complexes using the ubiquitin-like domain from HHR23B reveals novel proteasome-associated proteins. Biochemistry 48, 2538–2549 2009.

36. Kleijnen, M. F. et al. Stability of the proteasome can be regulated allosterically through engagement of its proteolytic active sites. Nat. Struct. Mol. Biol. 14, 1180–1188 2007.

37. Haselbach, D. et al. Long-range allosteric regulation of the human 26S proteasome by 20S proteasome-targeting cancer drugs. Nat. Commun. 8, 2017.

## References of methods

38. Farnung, L., Vos, S.M. & Cramer, P. Structure of transcribing RNA polymerase II-nucleosome complex. Nat Commun 9, 5432 (2018).

39. Wagner, T., Merino, F., Stabrin, M., Moriya, T., Antoni, C., Apelbaum, A., Hagel, P., Sitsel, O., Raisch, T., Prumbaum, D., Quentin, D., Roderer, D., Tacke, S., Siebolds, B., Schubert, E., Shaikh, T.R., Lill, P., Gatsogiannis, C. & Raunser, S. SPHIRE-crYOLO is a fast and accurate fully automated particle picker for cryo-EM. Communications Biology, 2, 218 (2019).

40. Zivanov, J., Nakane, T., Forsberg, B., Kimanius, D., Hagen, W. J. H., Lindahl, E., & Scheres, S.H.W. New tools for automated high-resolution cryo-EM structure determination in RELION-3. eLife, 7: e42166 (2018).

41. Steimle, S., Bajzath, C., Dörner, K., Schulte, M., Bothe, V. and Friedrich, T. Role of subunit NuoL for proton translocation by respiratory complex I. Biochemistry 50, 3386–3393 (2011).

42. Weissmann, F., Petzold, G., VanderLinden, R., Huis in ’t Veld, P.J., Brown, N.G., Lampert, F., Westermann, S., Stark, H., Schulman, B.A., Peters J.M. biGBac enables rapid gene assembly for the expression of large multisubunit protein complexes Proc. Natl. Acad. Sci. USA, 113, E2564–E2569 (2016).

43. Andersen, K.R., Leksa, N.C. & Schwartz, T.U. Optimized E. coli expression strain LOBSTR eliminates common contaminants from His-tag purification. Proteins 81, 1857–1861 (2013).

44. Kelley, K., Knockenhauer K.E., Kabachinski G. & Schwartz T.U., Atomic structure of the Y complex of the nuclear pore. Nature Structural & Molecular Biology 22, 425–431 (2015).

45. Besche, H. C., & Goldberg, A. L. Affinity Purification of Mammalian 26S Proteasomes Using an Ubiquitin-Like Domain. In R. J. Dohmen & M. Scheffner (Ed.), Ubiquitin Family Modifiers and the Proteasome (Vol. 832, pp. 423–432). Springer Science+Business Media, LCC. (2012).

