## Supplementary Information for "Quantifying the heterogeneity of macromolecular machines by mass photometry"

**Supplementary Movie 1:** First 5 seconds (real time) of a representative ratiometric movie of Complex I at 12.5 nM binding non-specifically to a glass coverslip. The field of view is 2.9×10.8 $\mu m^2$ and the raw frames were saved at 200 Hz, while the sliding ratiometric processing was applied with a frame summing of 5 frames. The movie is played back at 10 Hz.

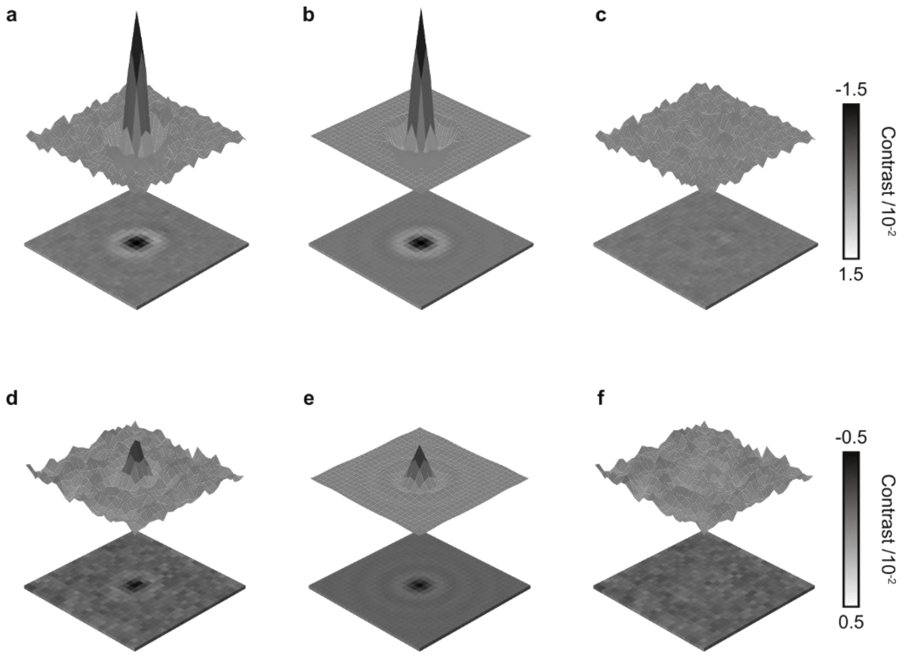


**Supplementary Figure 1:** Point spread function (PSF) fitting for mass photometry. Experimental (**a**, **d**) and fitted (**b**, **e**) PSF and the corresponding residual (**c**, **f**) for two particles with large (**a**, **b**, **c**) and small (**d**, **e**, **f**) signal-to-noise (SNR) ratios, SNR=12 and 3.5, respectively.

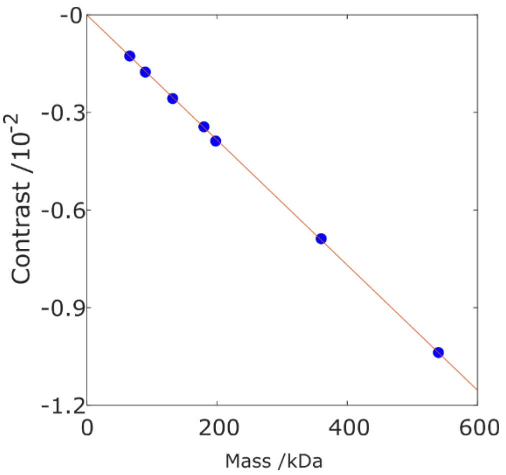


**Supplementary Figure 2:** Contrast-to-mass (C2M) calibration curve. The contrast of two known proteins with different oligomeric states were plotted vs their known mass. The red line is the fit to the data according to $y = bx$, with $b$ – C2M calibration factor.

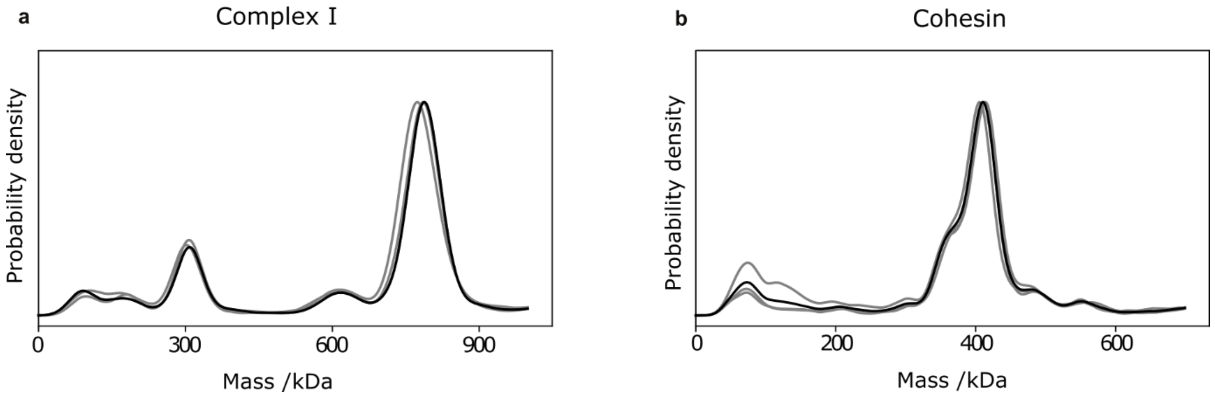


**Supplementary Figure 3:** Kernel density estimates (KDE) for repeats of mass photometry measurements for Complex I (**a**) and Cohesin (**b**) showing the reproducibility of MP for those complexes. Grey lines correspond to probability densities of 3 different independent measurements and black line is the combined probability density calculated from all particles. Number of particles of each repeat different are $N_{\mathrm{complex\,I}} = 2024, 2177, 1801$ and $N_{\mathrm{cohesin}} = 2196, 2031, 2046$.

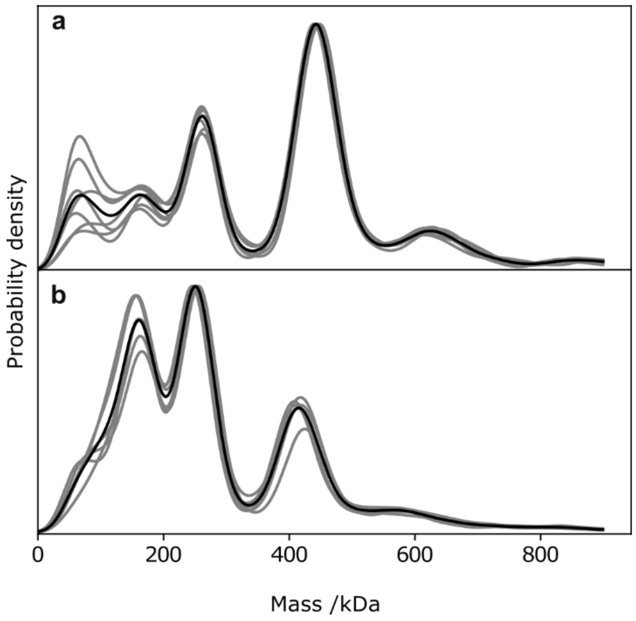


**Supplementary Figure 4:** Reproducibility of MP measurements for Nuclear Pore complex (NPC) before and after cross-linking. Probability density plots of MP for cross-linked NPC (**a**) for 5 min and without cross-linking treatment (**b**). Grey lines depict the probability density of 7 different independent measurements and the black line is the combined probability density.

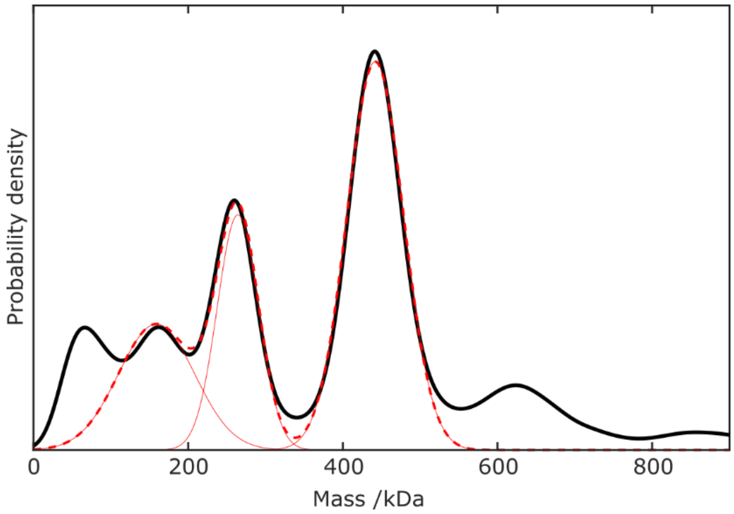


**Supplementary Figure 5:** Multiple Gaussian fitting to MP distributions of NPC. Black line corresponds to the experimental KDE, and red solid (dash) lines correspond to the 3 individual (sum of) fitted Gaussians.

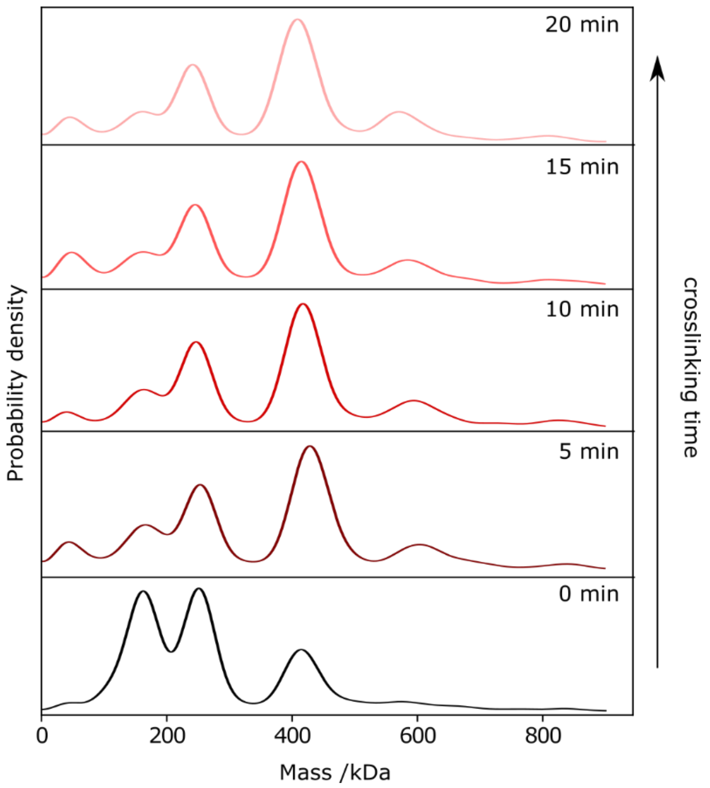


**Supplementary Figure 6:** Time dependent cross-linking of Nuclear Pore complex (NPC). MP probability density plots of NPC without any treatment (0 min), and after treatment with 0.1% glutaraldehyde for 5, 10, 15, 20 min, followed by quenching.

**Supplementary Table 1.** Molecular weights of individual APC/C subunits and APC/C subcomplexes. Subunits that form dimers are indicated with an asterisk.

| APC/C (1175.5 kDa) | | | | |
|---|---|---|---|---|
| **Subunit** | **Molecular weight (kDa)** | **Subunit** | **Molecular weight (kDa)** | |
| APC1 | 216 | APC8* | 69 | "Platform" subunits |
| APC2 | 94 | APC11 | 10 | |
| APC4 | 96 | APC13 | 9 | |
| APC5 | 85 | APC15 | 15 | |
| APC3* | 92 | APC10 | 21 | "Arc lamp" subunits |
| APC6* | 72 | APC12* | 10 | |
| APC7* | 67 | APC16 | 12 | |
| $APC/C^{Platform-APC8}$: 523 kDa | | | | |
| $APC/C^{Platform}$: 661 kDa | | | | |
| $APC/C^{Platform+APC6}$: 825 kDa | | | | |

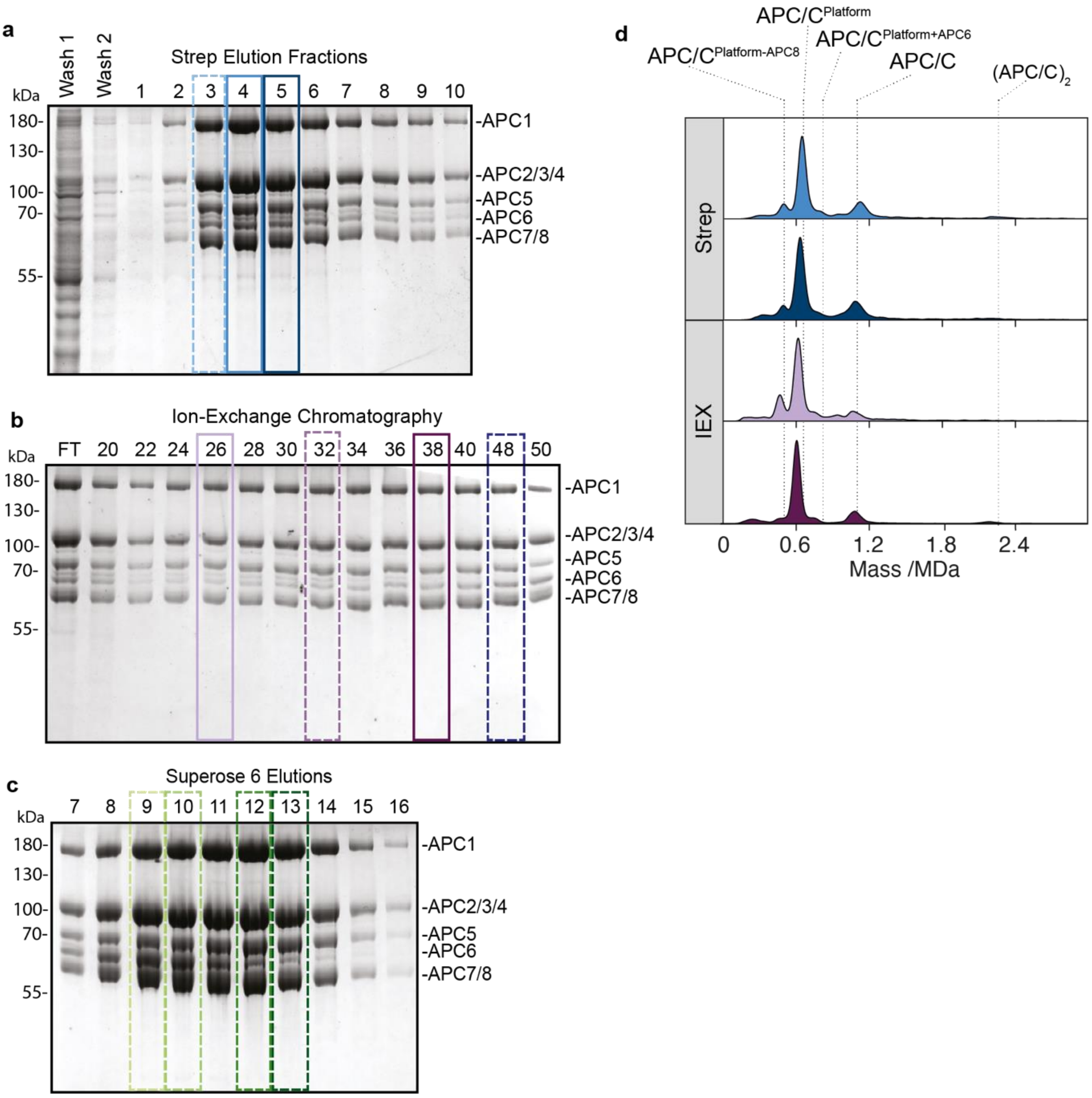


**Supplementary Figure 7. SDS PAGE and MP analysis of APC/C. a-c,** SDS PAGE analysis of APC/C purification steps. Boxed lanes indicate samples measured by MP as depicted in **d**. Dashed boxes indicate samples described in **Figure 2d-g.** Fractions 1-10 eluted from Strep-tactin (Strep) were combined for subsequent ion-exchange chromatography (IEX). Fractions 32-50 were combined for subsequent size-exclusion chromatography (SEC). **d,** MP measurements of fractions indicated in **a,b**.

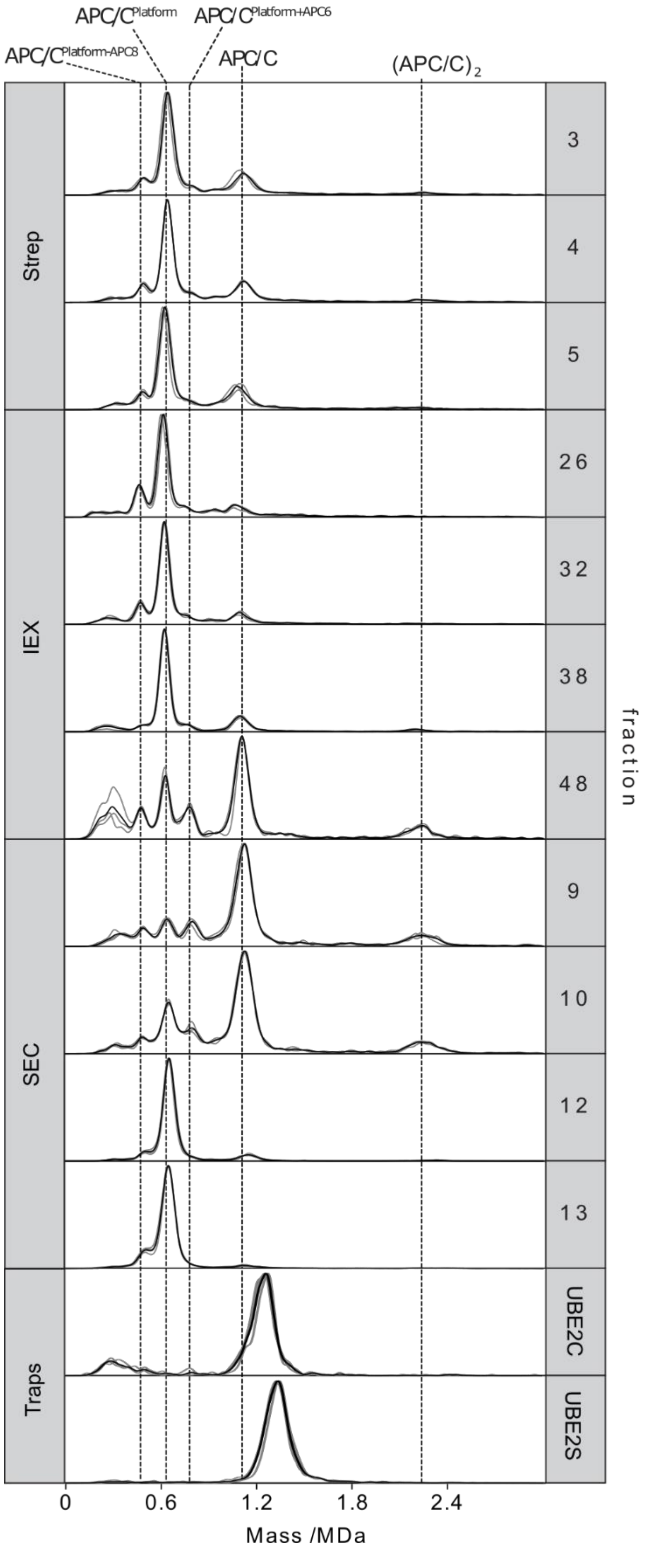


**Supplementary Figure 8:** Reproducibility of MP measurements for APC/C at its different purification steps, and two purified and cross-linked samples. Probability density plots of MP for different fractions of the three purification steps (Strep, IEX and SEC), and the two cross-linked APC/C: APC/C$^{CDH1}$-UBE2C and APC/C$^{CDH1}$-UBE2S traps indicated as UBE2C and UBE2S, respectively. Grey lines depict probability density of 3 different independent measurements and the black line represents the averaged probability density.

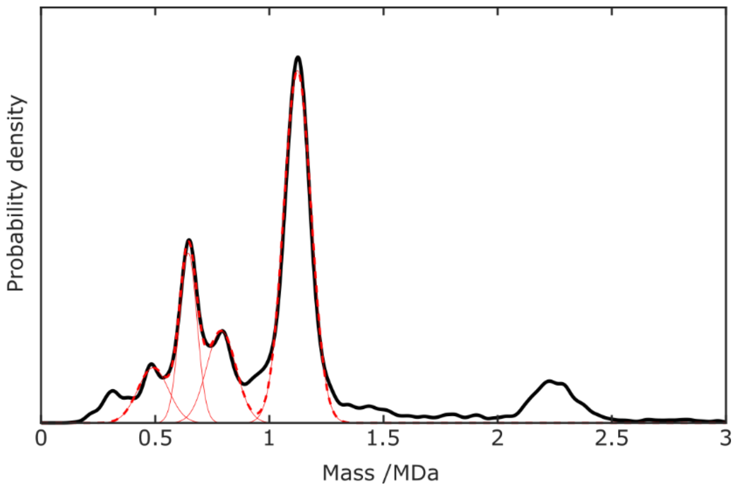


**Supplementary Figure 9:** Gaussian fitting to MP plots of APC/C. Four Gaussian fit to the KDE of SEC-fraction 10. The black line is the experimental KDE, and the red solid lines 4 individual fitted Gaussians, with the dashed lines representing their sum.

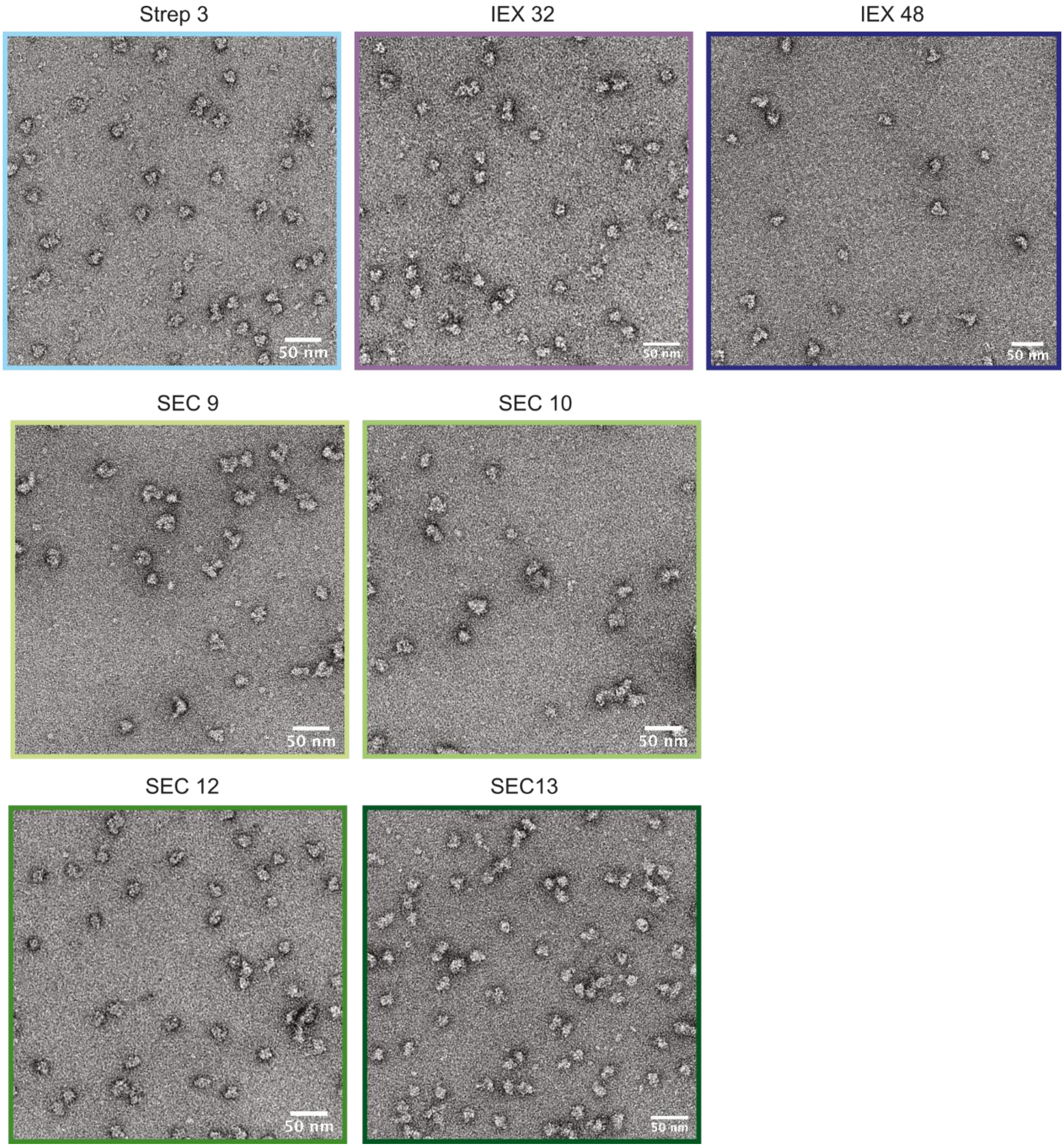


**Supplementary Figure 10:** Representative negative stain electron micrographs of APC/C fractions from Strep-tactin (Strep), ion-exchange chromatography (IEX), and size-exclusion chromatography (SEC) purification steps corresponding to 2D classifications depicted in **Figure 2g**. Scale bar: 50 nm.

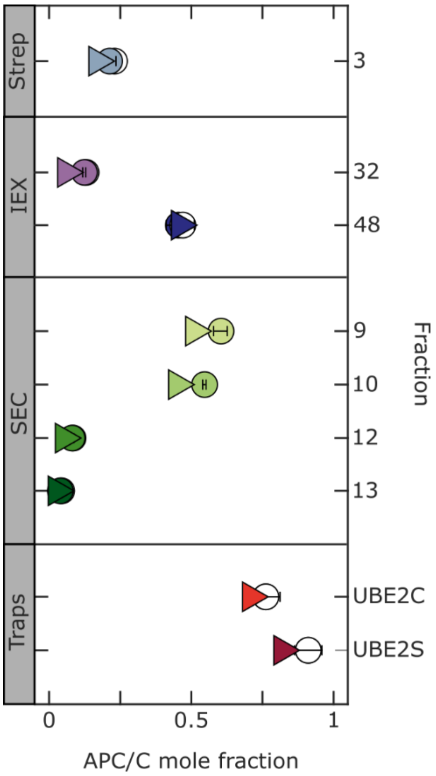


**Supplementary Figure 11:** Raw and diffusion-corrected mole fraction of APC/C obtained by MP, compared to nsEM. Mole fraction of full APC/C at the different steps of the purification steps (Strep, IEX and SEC) as well as for the two APC/C traps (APC/C cross-linked with cofactors) measured by MP (circles) and nsEM (triangles). 'Raw' MP mole fraction (empty circles) were corrected to account for the 'dead-time' (filled circles), resulting in a decrease in full APC/C fraction.

| | Strep | | | IEX | | | | SEC | | | | Traps | |
|---|---|---|---|---|---|---|---|---|---|---|---|---|---|
| | 3 | 4 | 5 | 26 | 32 | 38 | 48 | 9 | 10 | 12 | 13 | UBE2C | UBE2S |
| MP | 0.23 | 0.22 | 0.24 | 0.13 | 0.13 | 0.17 | 0.47 | 0.63 | 0.56 | 0.084 | 0.044 | 0.75 | 0.89 |
| MP corrected | 0.21 | 0.20 | 0.22 | 0.13 | 0.12 | 0.16 | 0.45 | 0.60 | 0.54 | 0.078 | 0.040 | - | - |
| nsEM | 0.17 | | | | 0.06 | | 0.46 | 0.51 | 0.45 | 0.052 | 0.027 | 0.71 | 0.82 |

**Supplementary Table 2:** APC/C Mole fraction at different steps of purification quantified by MP and nsEM. MP mole fractions were corrected for diffusion.

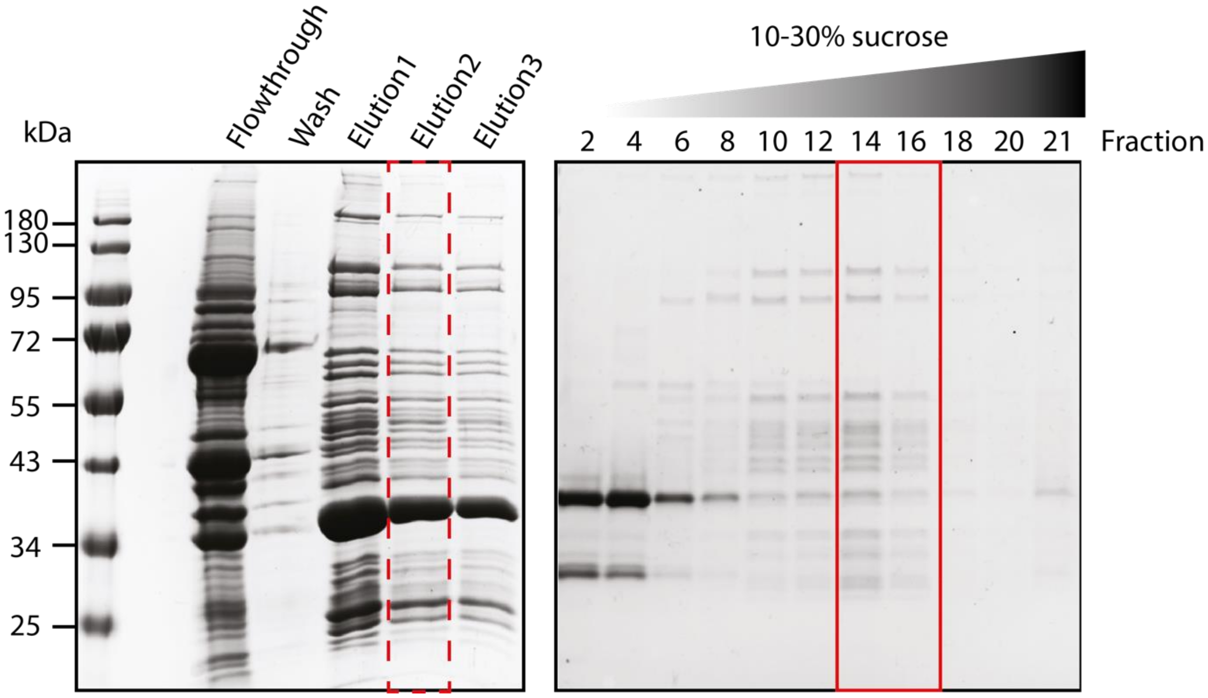


**Supplementary Figure 12**. SDS PAGE analysis of affinity purification of proteasome complexes from bovine heart. Elution2 is shown in **Fig. 3a**. Elution samples were applied on a 10-30% sucrose gradient. The boxed gradient fractions (14, 15 and 16) were pooled and used for MP and nsEM measurements.

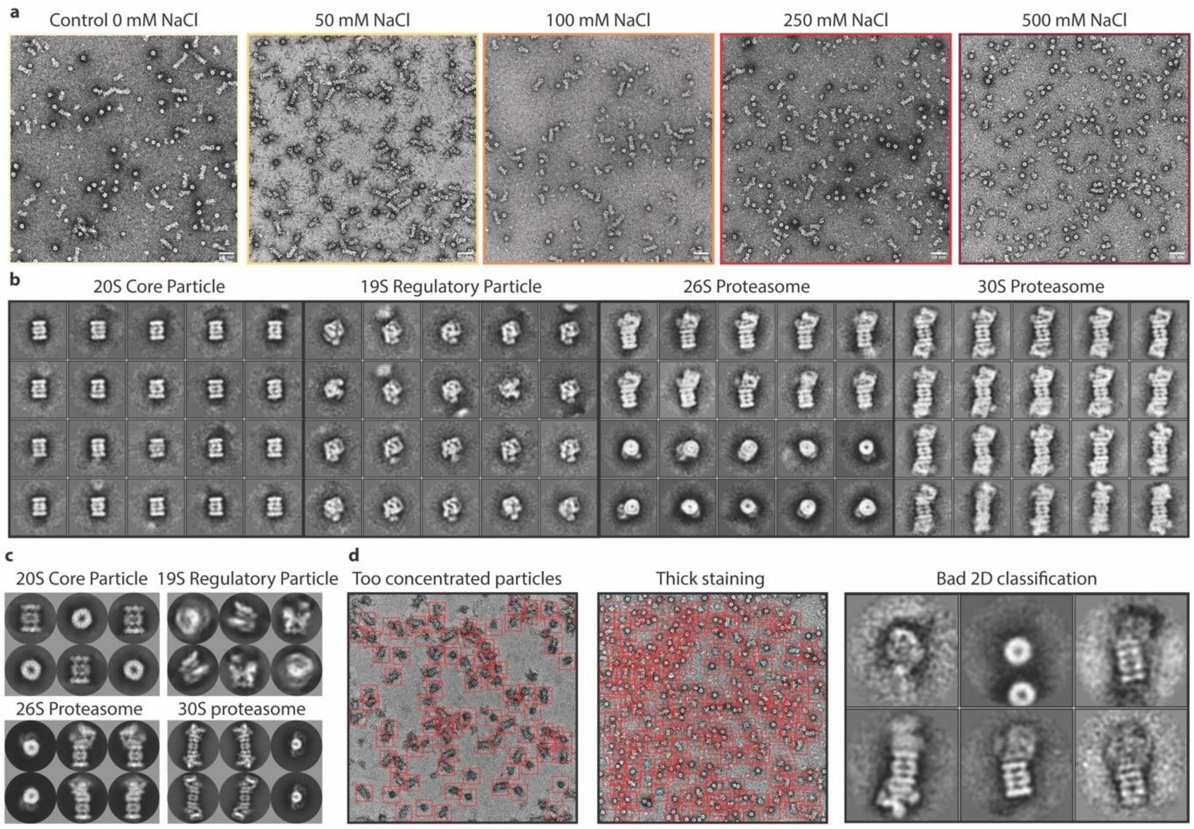


**Supplementary Figure 13:** Negative stain transmission electron microscopy analysis of proteasome samples. **a,** Representative negative stain electron micrographs for each salt concentration. Scale bar: 50 nm. **b,** Representative experimental 2D class averages of all proteasome complexes and subcomplexes. **c,** 2D projections (sampling at 120°) showing different views of proteasome complexes and subcomplexes. **d,** Examples of pitfalls in negative stain transmission electron microscopy sample preparation and data processing.

**Supplementary Table 3:** Molecular weights of individual proteasomal subunits and proteasome complexes and subcomplexes

| **20S core particle: 760 kDa** | | | |
|---|---|---|---|
| **Subunit** | **Molecular weight (kDa)** | **Subunit** | **Molecular weight (kDa)** |
| Proteasome subunit alpha-type 1 | 29.57 | Proteasome subunit beta-type 1 | 26.23 |
| Proteasome subunit alpha-type 2 | 25.88 | Proteasome subunit alpha-type 2 | 22.88 |
| Proteasome subunit alpha-type 3 | 28.39 | Proteasome subunit alpha-type 3 | 22.98 |
| Proteasome subunit alpha-type 4 | 29.47 | Proteasome subunit alpha-type 4 | 29.01 |
| Proteasome subunit alpha-type 5 | 26.39 | Proteasome subunit alpha-type 5 | 28.59 |
| Proteasome subunit alpha-type 6 | 27.38 | Proteasome subunit alpha-type 6 | 25.53 |
| Proteasome subunit alpha-type 7 | 27.85 | Proteasome subunit alpha-type 7 | 30.009 |
| **19S regulatory particle: 893 kDa** | | | |
| **Subunit** | **Molecular weight (kDa)** | **Subunit** | **Molecular weight (kDa)** |
| 26S proteasome regulatory subunit 7 (Rpt1) | 48.60 | 26S proteasome non-ATPase regulatory subunit 12 (Rpn5) | 60.92 |
| 26S proteasome regulatory subunit 4 (Rpt2) | 49.185 | 26S proteasome non-ATPase regulatory subunit 11 (Rpn6) | 53.02 |
| 26S proteasome regulatory subunit 6B (Rpt3) | 47.34 | 26S proteasome non-ATPase regulatory subunit 6 (Rpn7) | 47.43 |
| 26S proteasome regulatory subunit 10 (Rpt4) | 44.05 | 26S proteasome non-ATPase regulatory subunit 7 (Rpn8) | 45.5 |
| 26S proteasome regulatory subunit 6A (Rpt5) | 49.204 | 26S proteasome non-ATPase regulatory subunit 13 (Rpn9) | 36.73 |
| 26S proteasome regulatory subunit 8 (Rpt6) | 45.6 | 26S proteasome non-ATPase regulatory subunit 4 (Rpn10) | 42.84 |
| 26S proteasome non-ATPase regulatory subunit 2 (Rpn1) | 100.19 | 26S proteasome non-ATPase regulatory subunit 14 (Rpn11) | 41.36 |
| 26S proteasome non-ATPase regulatory subunit 1 (Rpn2) | 105.836 | 26S proteasome non-ATPase regulatory subunit 8 (Rpn12) | 34.57 |
| 26S proteasome non-ATPase regulatory subunit 3 (Rpn3) | 60.92 | 26S proteasome non-ATPase regulatory subunit 15 (Sem1) | 32.52 |
| **20S core particle – 19S regulatory particle: 1653 kDa** | | | |
| **19S regulatory particle - 20S core particle – 19S regulatory particle: 2546 kDa** | | | |

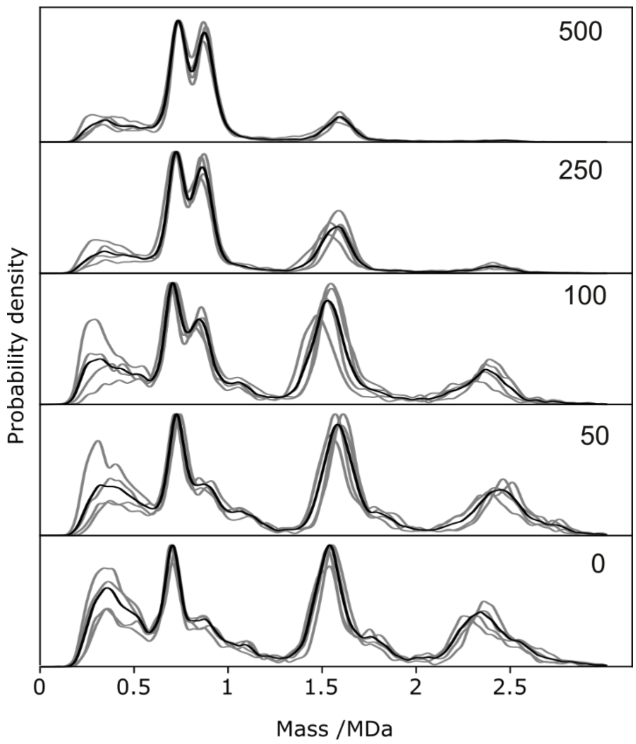


**Supplementary Figure 14:** Reproducibility of MP measurements for proteasome at different NaCl concentrations. Probability density plots of MP for different NaCl concentrations in mM. Grey lines correspond to the probability density of 4 different independent measurements (and at two different days) and the black line is the combined probability density.

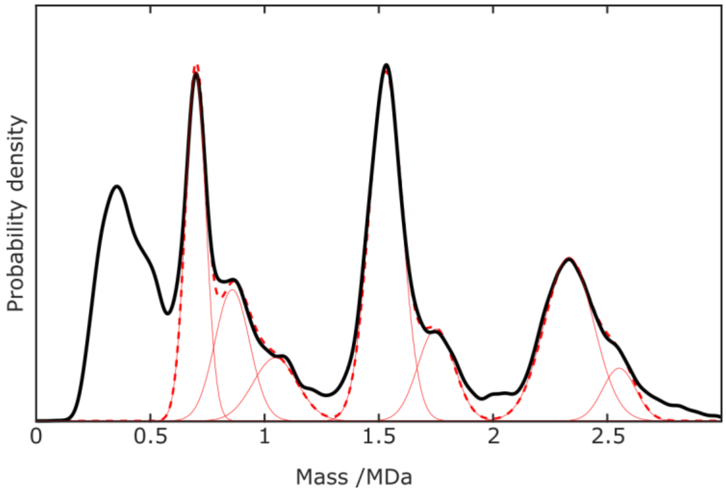

**Supplementary Figure 15:** Gaussian fitting to MP distribution for the proteasome. Seven Gaussian fitting to the KDE at 0 mM NaCl. Black line corresponds to the experimental KDE, and red solid (dash) lines correspond to the 7 individual (sum of) fitted Gaussians.

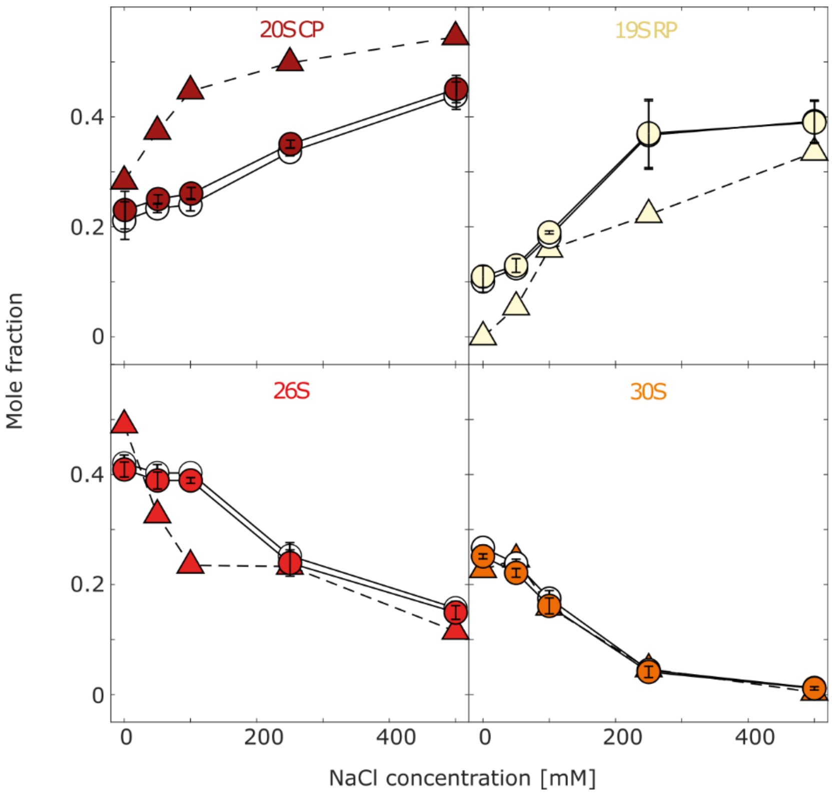


**Supplementary Figure 16:** Diffusion corrected mole fraction for proteasome treatment with salt. Mole fraction vs NaCl concentration for the 4 different proteasome (sub-) complexes, 20S, 19S RP, 26S and 30S, measured by MP (circles) and nsEM (triangles). 'Raw' MP mole fraction (empty circles) were corrected to account for the 'dead-time' (filled circles), resulting in an increase of the mole fractions for small species.

| Salt | 20S | | | 19S | | | 26S | | | 30S | | |
|---|---|---|---|---|---|---|---|---|---|---|---|---|
| | MP | MP corrected | nsEM | MP | MP corrected | nsEM | MP | MP corrected | nsEM | MP | MP corrected | nsEM |
| 0 | 0.21 | 0.23 | 0.28 | 0.10 | 0.11 | 0 | 0.42 | 0.41 | 0.49 | 0.27 | 0.25 | 0.23 |
| 50 | 0.24 | 0.25 | 0.37 | 0.12 | 0.13 | 0.054 | 0.40 | 0.39 | 0.33 | 0.24 | 0.22 | 0.25 |
| 100 | 0.24 | 0.26 | 0.44 | 0.18 | 0.19 | 0.16 | 0.40 | 0.39 | 0.23 | 0.17 | 0.16 | 0.16 |
| 250 | 0.34 | 0.35 | 0.50 | 0.37 | 0.37 | 0.22 | 0.25 | 0.24 | 0.23 | 0.045 | 0.041 | 0.046 |
| 500 | 0.44 | 0.45 | 0.54 | 0.39 | 0.39 | 0.34 | 0.16 | 0.15 | 0.11 | 0.012 | 0.011 | 0.004 |

**Supplementary Table 4:** Proteasome mole fraction analysis for its different (sub-) complexes, at increasing salt concentration, quantify by MP and nsEM. MP mole fractions were corrected for the different diffusion of (sub-) complex, and are in good agreement with nsEM ones.

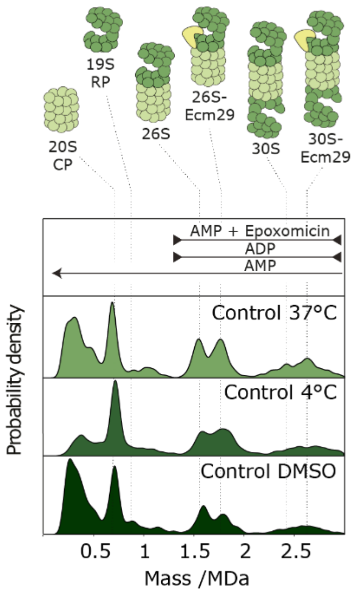


**Supplementary Figure 17:** MP control experiments for proteasome samples at 4°C, 37°C and in the presence of DMSO, for the proteasome sample used for different nucleotide conditions (**Fig. 3e** of main text). All measurements were performed at 50 nM.

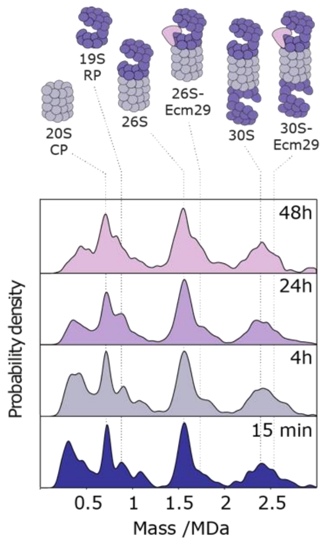


**Supplementary Figure 18:** MP measurements of proteasome stability. Probability density of proteasome at different time points (15 min, 4h, 24h and 48h) show that proteasome (sub)complexes are stable even after 48 hours. All samples were kept at 4°C and measured at 50 nM.

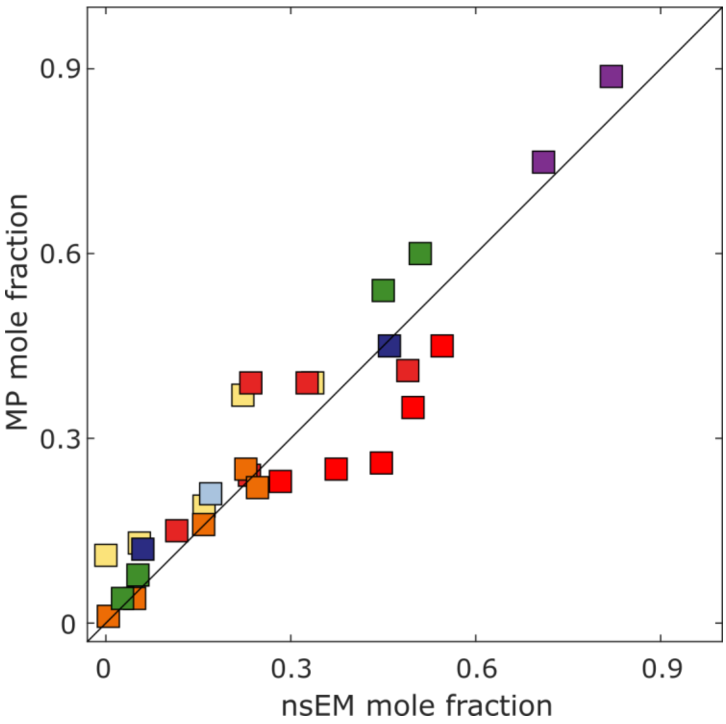


**Supplementary Figure 19:** Mass photometry and negative staining comparison. Scatter plot of mole fraction in MP vs nsEM for all analyzed samples of APC/C purification steps and Proteasome salt addition. Colors correspond to different protein samples: APC/C in purification steps: strep (light-blue), IEX (dark-blue), SEC (green), APC/C cross-linked (purple); Proteasome – 20S (brown), 19S (yellow), 26S (red) and 30S (orange). Solid line corresponds to $y = x$, with $R^2$=0.9.
